# The IFN-γ-inducible GTPase IIGP1 regulates mitochondrial remodeling and immunometabolic reprogramming during inflammatory macrophage polarization

**DOI:** 10.64898/2026.11.02.763539

**Authors:** Anup Kumar Singh, Samriddhi Sharma, Kriti Ahuja, Rajender K Motiani, Srikanth Rapole, Manas Kumar Santra, Arvind Sahu

## Abstract

Macrophages are key effector cells of the innate immune system that undergo dynamic polarization into functionally distinct phenotypic states, ranging from classically activated inflammatory (M1) to alternatively activated (M2) macrophages. Inflammatory macrophage polarization requires coordinated integration of cytokine signaling, metabolic reprogramming, and mitochondrial remodeling. However, the molecular effectors that control inflammatory signaling for mitochondrial metabolism and remodeling remain poorly defined. Using quantitative mass spectrometry-based proteomics, we identified Interferon-inducible GTPase 1 (IIGP1) as one of the most robustly induced proteins during inflammatory macrophage polarization. Its expression is governed by the IFN-γ–JAK-STAT1 signaling axis and is crucial for this polarization process. IIGP1-deficient macrophages exhibited significantly impaired iNOS induction, nitric oxide production, pro-inflammatory gene expression, co-stimulatory surface marker expression, and morphological remodeling. Further, IIGP1 associates with mitochondria and its absence impairs key characteristics of inflammatory macrophages, including glycolytic metabolic reprogramming, mitochondrial redox adaptation, membrane potential regulation, and structural remodeling. Consistent with these defects, IIGP1-deficient macrophages displayed impaired intracellular killing of Escherichia coli and Salmonella Typhimurium. Collectively, these results point to IIGP1 as an IFN-γ-inducible effector that facilitates efficient inflammatory macrophage polarization and antibacterial immunity by linking inflammatory signaling to mitochondrial homeostasis and immunometabolic reprogramming.

## Introduction

Macrophages are the key effector cells of the innate immune system that play a critical role in host defense, inflammation, tissue homeostasis, and repair (1). These highly versatile cells coordinate immune responses against invasive pathogens and shape the subsequent adaptive immune responses by carrying out phagocytosis, antigen presentation, cytokine release, and the generation of antimicrobial and immunomodulatory mediators like nitric oxide (NO) and reactive oxygen species (ROS) (2, 3, 4). Additionally, macrophages have a remarkable ability of functional plasticity, which allows them to respond dynamically to their environmental signals and adopt a unique polarized state depending on the physiological and pathological conditions (5).

Macrophage polarization is a highly dynamic and plastic process involving a continuum of phenotypic states ranging from the classically activated inflammatory (M1) macrophages to alternatively activated (M2) macrophages (6). M1 polarization is driven by type 1 cytokine signals, including IFN-γ and TNF-α, as well as microbial components such as LPS. This results in the production of pro-inflammatory cytokines like IL-6, IL-1β, and TNF-α, with reactive oxygen/nitrogen species (ROS/RNS) and enhanced antimicrobial activity (2, 6). Conversely, type 2 cytokine signals like interleukin-4 (IL-4) and/or IL-13 drive the development of the M2 macrophage phenotype, which is associated with the secretion of anti-inflammatory cytokines like IL-10 and TGF-β and is linked to tissue remodeling and the resolution of inflammation (6–9).

Distinct metabolic states also play a vital role in characterization of these different functional macrophage phenotypes (10). Pro-inflammatory M1 phenotypes are marked by enhanced aerobic glycolysis (11–12), whereas anti-inflammatory M2 phenotypes predominantly rely on oxidative phosphorylation and fatty acid oxidation (13). Notably, these metabolic differences involve mitochondrial remodeling, which leads to changes in mitochondrial activity, redox homeostasis, and altered morphology and dynamics (14–16), thereby highlighting the vital role of mitochondria in the regulation of macrophage polarization and their effector function.

Growing evidences suggest that a coordinated integration of cytokine signaling, mitochondrial remodeling, and metabolic reprogramming is required for the inflammatory macrophage polarization (17–19). Beyond their canonical role in energy production, mitochondria serve as essential signaling hubs that regulate the generation of ROS, inflammatory signaling pathways, immune responses, and antimicrobial effector mechanisms. Consistent with this, disruption of mitochondrial homeostasis has been shown to modulate macrophage polarization and immune function (20–22).

IFN-γ signaling plays a central role in driving inflammatory macrophage polarization. Activation of its downstream JAK-STAT1 pathway induces a broad transcriptional program that promotes antimicrobial and inflammatory functions, characteristic of M1 macrophages (25). Despite extensive characterization of IFN-γ-dependent transcriptional programs, the downstream effector proteins that are required for acquisition and maintenance of the inflammatory macrophage phenotype remain incompletely understood. Among the proteins induced downstream of IFN-γ-signaling, interferon-responsive GTPases are particularly strong candidates because of their established roles in host defense and intracellular membrane remodeling. The immunity-related GTPases (IRGs) constitute a family of IFN-γ inducible proteins belonging to the dynamin-GTPase superfamily (26) and function as important mediators of host defense against a wide range of intracellular pathogens (27–35). Emerging evidence indicates that IRG family proteins such as IRGM, a human IRG family member that associates with mitochondria and regulates mitochondrial dynamics, can also regulate mitochondrial biology (36) and mitochondrial remodeling and autophagy-related processes (37), suggesting that mitochondrial regulation may be a conserved feature of IRG family proteins.

IRG proteins participate in diverse cellular processes, including autophagy, intracellular membrane remodeling, vesicular trafficking, and cytoskeletal regulation (29, 34, 38–40). p47 IRG family (41) members are well-established regulators of cell-autonomous immunity, and IIGP1, which belongs to this family, is a murine interferon-inducible GTPase. IIGP1 has been shown to regulate cell motility (39), contribute to autophagy (42), and facilitate recruitment of non-IRG effector proteins, including guanylate-binding proteins (GBPs) and Atg8/LC3-associated autophagy machinery, to pathogen-containing vacuoles (38). Thus, IIGP1 contributes to remodeling of membrane-associated compartments that are critical for host resistance against intracellular pathogens. IIGP1 has been shown to localize predominantly to intracellular membrane compartments, such as the Golgi apparatus and endoplasmic reticulum, where it participates in intracellular membrane organization and antimicrobial responses (42). In addition to these compartments, IIGP1 has also been shown to associate with mitochondria, suggesting a role in organelle-specific regulatory processes (43). Given the central role of mitochondrial remodeling in sustaining inflammatory macrophages metabolism, suggesting that IIGP1 may function as a previously unrecognized regulator of mitochondrial homeostasis during inflammatory macrophage polarization. However, whether IIGP1 contributes to mitochondrial organization and metabolic adaptation during inflammatory macrophage polarization has not been investigated.

Although major signaling pathways and transcriptional regulators governing macrophage polarization have been extensively characterized (23, 24), the molecular effectors that control inflammatory signaling to regulate mitochondrial remodeling and metabolic adaptation remain poorly understood. In particular, how coordinated mitochondrial remodeling and redox adaptation are sustained during inflammatory macrophage activation and which proteins mediate this process have not been established (20–22). In this study, we address this gap by identifying IIGP1, through whole-proteome analysis of IFN-γ-polarized macrophages, as an IFN-γ-inducible effector that connects inflammatory signaling to mitochondrial homeostasis and immunometabolic reprogramming in macrophages. IIGP1 (encoded by Irga6 in mice) is one of the most robustly induced proteins during inflammatory macrophage activation.

Further, we investigated the role of IIGP1 in IFN-γ-driven macrophage polarization and immunometabolic remodeling. We demonstrated that IIGP1 functions downstream of IFN-γ-JAK-STAT1 signaling and is required for the establishment of the inflammatory macrophage phenotype. Mechanistically, IIGP1 associates with mitochondria and supports the glycolytic metabolic reprogramming, redox regulation, and mitochondrial remodeling characteristic of inflammatory macrophages. Loss of IIGP1 disrupts mitochondrial organization and metabolic remodeling, impairs inflammatory macrophage polarization, and compromises antibacterial effector function. Collectively, our findings identify IIGP1 as a previously unrecognized IFN-γ-inducible effector that links inflammatory signaling to mitochondrial homeostasis, metabolic adaptation, and antibacterial effector function in macrophages.

## Results

### Proteomics profiling identifies IIGP1 as a selectively enriched protein during inflammatory macrophage polarization

To identify downstream effector proteins associated with macrophage polarization, we performed mass spectrometry-based global proteomic profiling of primary bone marrow-derived macrophages (BMDMs) and RAW264.7 cells under unstimulated (M0), IFN-γ-stimulated inflammatory (M1), and IL-4-stimulated alternative (M2) activation conditions (**Fig. 1A**). Successful macrophage polarization was initially confirmed through detailed morphological characterization and the validation of canonical activation markers. Polarized macrophages exhibited distinct structural remodeling, with phenotype-specific morphological changes clearly evident in both BMDMs (**Fig. 1B**; quantified in **Fig. S1A and C)** and RAW264.7 cells (**Fig. 1C**; quantified in **Fig. S1D and F**). At the transcriptional level, quantitative RT-PCR analysis of lineage-specific markers, including *Nos2*, *Il-6*, and *Il-1β* for the M1 phenotype, along with *Arg1*, *Mrc1*, *Ym1*, and *Fizz1* for the M2 phenotype, confirmed robust polarization in BMDMs **(Fig. S2A and B)**. Similarly, successful transcriptional reprogramming was validated in RAW264.7 cells via the induction of *Nos2*, *Il-6*, and *Il-1β* under M1 conditions and *Arg1* and *Mrc1* under M2 conditions **(Fig. S2C and D)**. Consistent with these transcriptomic shifts, immunoblot analysis demonstrated a robust iNOS protein expression in M1 macrophages and Arg-1 expression in M2 macrophages across both cell models **(Fig. S2E and F)**, confirming successful polarization relative to the baseline M0 phenotype.

**Figure 1.**
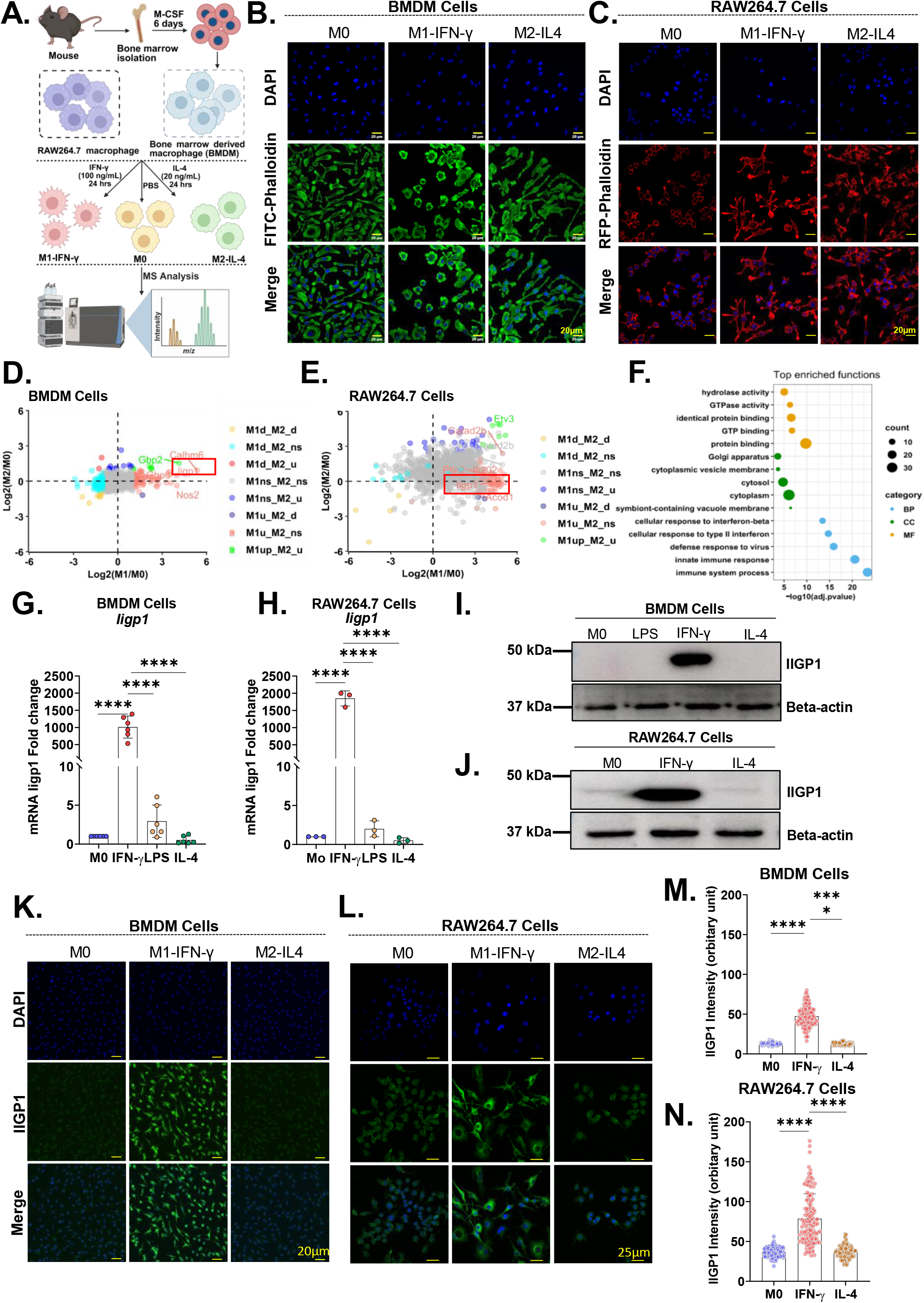
Proteomic profiling identifies IIGP1 as a selectively enriched protein during inflammatory macrophage polarization. **(A)** Schematic representation of bone marrow-derived macrophage (BMDM) generation, RAW264.7 cell culture, and polarization workflows. Cells were stimulated for 24 h with IFN-γ (100 ng/mL) to induce M1 polarization or with IL-4 (10 ng/mL for BMDMs; 20 ng/mL for RAW264.7 cells) to induce M2 polarization. **(B, C)** Representative immunofluorescence images showing the morphology of M0, M1, and M2 macrophages. Cells were counterstained with DAPI to visualize nuclei (blue) along with **(B)** Alexa Fluor™ 488-phalloidin (green) to visualize F-actin in BMDMs or **(C)** Rhodamine-phalloidin (red) to visualize F-actin in RAW264.7 cells. **(D, E)** Volcano plots illustrating global quantitative proteomic profiles of polarized macrophages relative to unstimulated (M0) controls in **(D)** BMDMs and **(E)** RAW264.7 cells. **(F)** Enriched Kyoto Encyclopedia of Genes and Genomes (KEGG) pathways and Gene Ontology (GO) biological processes derived from differentially expressed proteins in M1 macrophages. **(G, H)** Quantitative RT-PCR analysis of *Iigp1* mRNA induction in **(G)** BMDMs and **(H)** RAW264.7 cells across polarization conditions; transcript levels are presented relative to M0 controls. **(I, J)** Immunoblot analysis of IIGP1 expression in whole-cell lysates from **(I)** BMDMs and **(J)** RAW264.7 cells, with β-actin serving as the loading control. **(K–N)** Representative immunofluorescence microscopy and corresponding signal intensity quantification for cellular IIGP1 expression. Polarized cells were fixed in 4% paraformaldehyde and probed with an anti-IIGP1 primary antibody followed by an Alexa Fluor™ 488-conjugated secondary antibody. Panels depict **(K)** representative micrographs in BMDMs, **(L)** representative micrographs in RAW264.7 cells, **(M)** fluorescence intensity quantification in BMDMs, and **(N)** fluorescence intensity quantification in RAW264.7 cells. Data represent mean ± SD (n=3 independent biological replicates). Statistical significance was determined by one-way ANOVA with Tukey’s post-hoc test (*p* ≤ is less than or equal to ≤ 0.05, ** *p* ≤ is less than or equal to ≤ 0.01, *** *p* ≤ is less than or equal to ≤ 0.001, **** *p* ≤ is less than or equal to ≤ 0.0001).

Comparative proteomic analysis revealed substantial remodeling of the macrophage proteome upon activation. Both BMDMs (**Fig. 1D**) and RAW264.7 cells (**Fig. 1E**) displayed highly distinct, stimulus-specific proteomic signatures following IFN-γ or IL-4 treatment relative to baseline M0 controls. Functional annotation and Gene Ontology (GO)-based enrichment analysis of differentially expressed proteins in M1-polarized RAW264.7 macrophages revealed a significant enrichment of pathways governing innate immune activation (**Fig. 1F**). Importantly, molecular function enrichment analysis within the M1 proteome identified a strong overrepresentation of GTP-binding and GTPase-associated proteins, suggesting the involvement of GTPase-mediated regulatory mechanisms during classical macrophage activation. Furthermore, GO cellular component analysis highlighted significant enrichment of proteins localized to the cytoplasm, Golgi apparatus, and vesicle membrane compartments (**Fig. 1F**).

Among the differentially expressed candidates, IIGP1 emerged as a compelling protein for further investigation due to its status as one of the most robustly induced proteins following IFN-γ-mediated inflammatory polarization. IIGP1 belongs to the immunity-related GTPase (IRG) family, whose members are well-established mediators of intracellular antimicrobial host defense. Given previous literature documenting its association with intracellular membrane compartments, including mitochondria, together with emerging evidence implicating IRG family proteins in organelle homeostatic control, we hypothesized that IIGP1 might act as a molecular effector linking upstream inflammatory signaling to mitochondrial remodeling during macrophage polarization.

To validate our proteomic findings, we first verified IIGP1 expression profiles using quantitative RT-PCR. IIGP1 transcript levels were significantly and selectively upregulated in M1 macrophages compared to M0 and M2 states in both BMDMs (**Fig. 1G**) and RAW264.7 cells (**Fig. 1H**). This trend was confirmed at the translational level by Western blot, which revealed a highly statistically significant increase in IIGP1 protein expression exclusively within the M1 phenotype across both BMDMs (**Fig. 1I**) and RAW264.7 cells (**Fig. 1J**). Additionally, immunofluorescence microscopy and corresponding signal intensity quantification in BMDMs (**Fig. 1K and M**) and RAW264.7 cells (**Fig. 1L and N**) confirmed robust, selective cytoplasmic expression of IIGP1 protein upon IFN-γ stimulation. Collectively, these findings establish IIGP1 as a polarization-dependent protein that is selectively enriched during IFN-γ-driven inflammatory macrophage polarization.

### IFN-γ-JAK-STAT1 signaling regulates IIGP1 expression during inflammatory macrophage polarization

Given the selective induction of IIGP1 in M1 macrophages, we examined the upstream signaling mechanism controlling IIGP1 expression during inflammatory activation. Initially, we performed STRING protein interaction network analysis and found a clear functional association between IIGP1 and STAT1, which is a central mediator of IFN-γ signaling (**Fig. 2A**). Further, in-silico promoter analysis showed a putative STAT1 binding motif (GAS elements) within the Iigp1 promoter region, suggesting a potential transcriptional regulation of IIGP1 by STAT1 (**Fig. 2B**).

**Figure 2.**
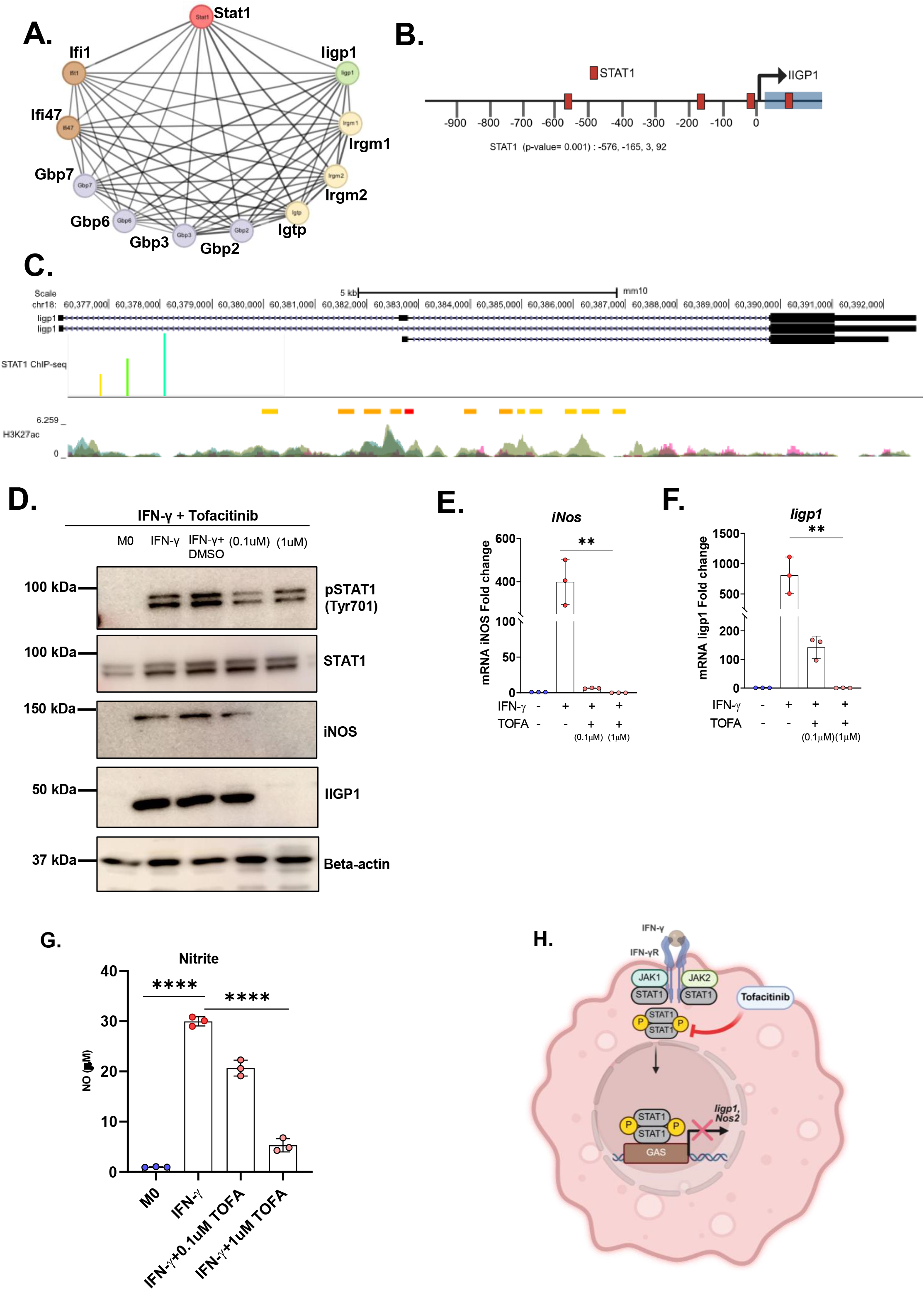
IFN-γ-JAK-STAT1 signaling regulates IIGP1 expression during inflammatory macrophage Polarization. **(A)** STRING database network analysis highlighting STAT1 as a core transcription factor associated with IIGP1. **(B)** In silico analysis mapping predicted canonical STAT1-binding GAS elements upstream of the mouse *Iigp1* transcription start site. **(C)** UCSC Genome Browser tracks (GSE33913) showing STAT1 recruitment and H3K27ac enrichment at the *Iigp1* locus in macrophages after IFN-γ stimulation. **(D)** Immunoblot of RAW264.7 lysates treated with IFN-γ (100 ng/mL) ±plus or minus ± tofacitinib (TOFA; 0.1-1 µM), probed for p-STAT1, STAT1, IIGP1, iNOS, and *β*-actin (loading control). **(E)** Quantitative RT-PCR (qPCR) of *iNOS* mRNA expression across treatment groups. **(F)** Quantitative RT-PCR (qPCR) of *Iigp1* mRNA expression across treatment groups. **(G)** Nitric oxide (NO) production in culture supernatants measured via Griess assay. **(H)** Schematic model of the IFN-γ-JAK-STAT1 axis driving *Iigp1* transcription, and its inhibition by TOFA. Data represent mean ± SD (n=3 independent biological replicates). Statistical significance was determined by one-way ANOVA with Tukey’s post-hoc test (*p* ≤ is less than or equal to ≤ 0.05, ** *p* ≤ is less than or equal to ≤ 0.01, *** *p* ≤ is less than or equal to ≤ 0.001, **** *p* ≤ is less than or equal to ≤ 0.0001).

To substantiate that STAT1 physically occupies these predicted regulatory sequences *in vivo*, we analyzed public functional genomics data tracking genomic STAT1 recruitment in bone marrow-derived macrophages (BMDMs) stimulated with IFN-γ for 6 hours (GEO: GSE33913). Chromatin immunoprecipitation sequencing (ChIP-seq) track visualization revealed striking, statistically significant STAT1 enrichment peaks localized directly over the *Iigp1* proximal promoter immediately upstream of the transcription start site, as well as within its first intron (**Fig. 2C**). These structural docking sites overlapped precisely with localized peaks of H3K27ac histone modifications, demonstrating that STAT1 actively targets transcriptionally receptive, open chromatin environments to orchestrate IIGP1 expression. To functionally validate the requirement of this JAK-STAT axis in driving IIGP1 induction, we stimulated macrophages with IFN-γ in the presence or absence of the JAK-STAT pathway inhibitor tofacitinib (TOFA). We confirmed that TOFA treatment successfully suppressed the IFN-γ-mediated phosphorylation of STAT1 (**Fig. 2D**). Consistent with a STAT1-dependent inflammatory program, IFN-γ stimulation significantly increased iNOS transcript expression and subsequent nitric oxide production, both of which were markedly blunted upon TOFA administration (**Fig. 2E and G**). Crucially, the downstream IFN-γ-stimulated induction of both IIGP1 transcripts (**Fig. 2F**) and translation into mature protein (**Fig. 2D**) was significantly abolished following TOFA treatment compared to the stimulated control group. Collectively, these findings provide definitive physical and functional evidence identifying the IFN-γ-JAK-STAT1 signaling cascade as the critical upstream regulatory pathway directly controlling IIGP1 expression during inflammatory macrophage polarization (**Fig. 2H**).

### IIGP1 is intrinsically required for IFN-γ-induced iNOS expression and nitric oxide production during inflammatory macrophage polarization

Preceding results demonstrate that IIGP1 is induced through JAK-STAT1 signaling pathway during IFN-γ-mediated M1 polarization. We next investigated whether it contributes to the acquisition and maintenance of the inflammatory macrophage phenotype. To address this, we generated IIGP1-deficient RAW264.7 macrophages using CRISPR-Cas9-mediated genome editing targeting exon 2 of the *Iigp1* locus (**Fig. 3A**). Successful gene ablation was validated by immunoblotting of whole-cell lysates (**Fig. 3B**) and immunofluorescence (**Fig. 3C**), which demonstrated a complete absence of IIGP1 protein in knockout cells following IFN-γ stimulation. To determine whether IIGP1 contributes to macrophage polarization, we analyzed expression of canonical M1 marker iNOS. Both immunoblotting and immunofluorescence staining revealed a significant reduction in iNOS expression in IIGP1-deficient M1 macrophages relative to wild-type (WT) controls (**Fig. 3D, E and F)**. Consistent with this down-regulation, IIGP1-deficient cells exhibited markedly impaired nitric oxide (NO) production under M1-polarizing conditions, whereas no altered baseline was observed in M0 or M2 states (**Fig. 3G**).

**Figure 3.**
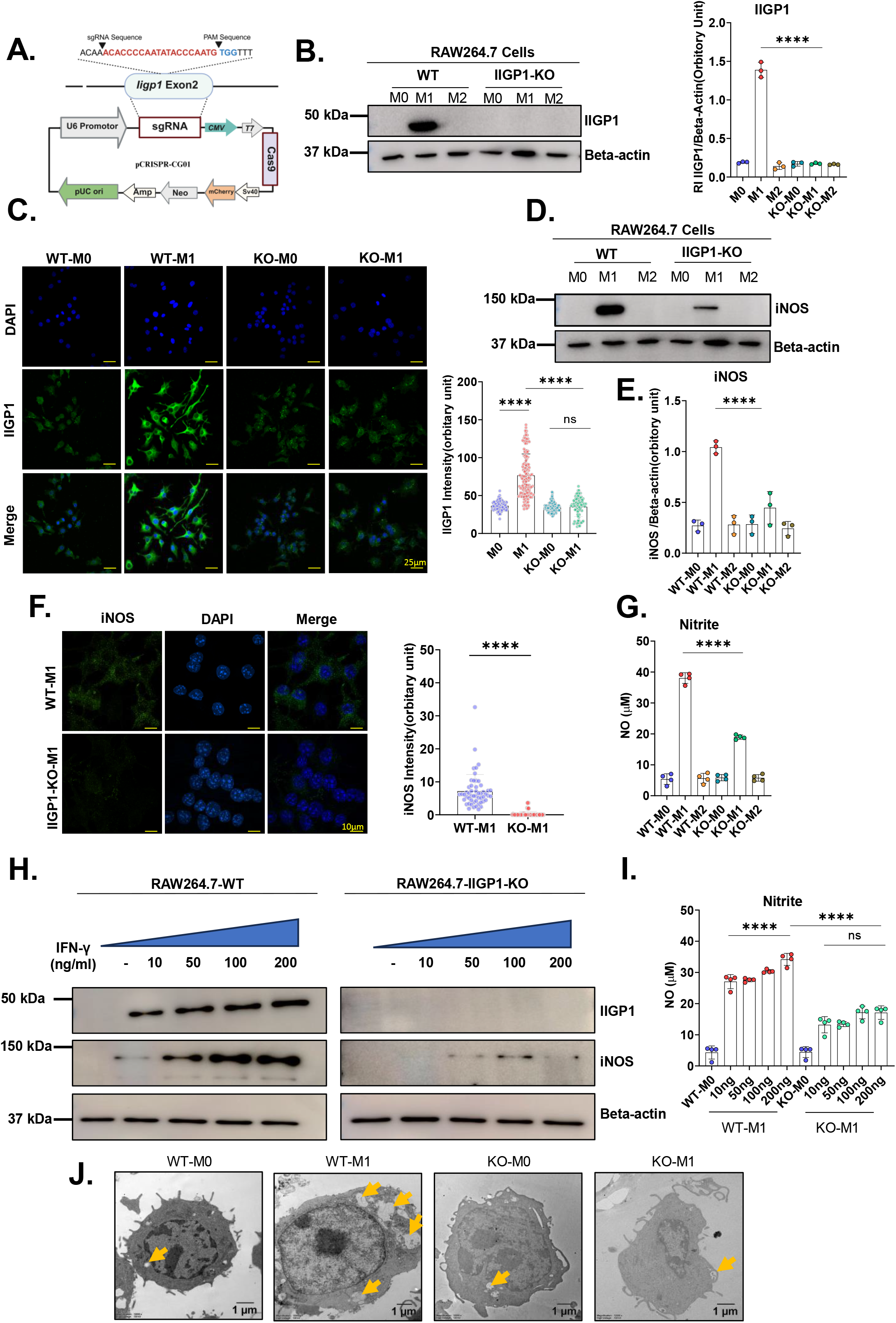
IIGP1 is intrinsically required for IFN-γ-induced iNOS expression and nitric oxide production during inflammatory macrophage polarization. **(A)** Schematic representation of the CRISPR-Cas9 strategy used to generate *Iigp1*-knockout (IIGP1-KO) RAW264.7 macrophages, including sgRNA sequences. **(B)** Immunoblot and quantification of **IIGP1** in WT and IIGP1-KO RAW264.7 lysates under M0, M1 (IFN-γ), and M2 **(IL-4)** conditions (24 h); *β*-actin served as loading control. **(C)** Immunofluorescence analysis and quantified fluorescence intensity of IIGP1 (green) in WT and IIGP1-KO macrophages under M0 and M1 conditions (24 h); nuclei counterstained with DAPI (blue). scale bar = 25 μm. **(D)** Immunoblot and quantification of iNOS expression in WT and IIGP1-KO cells under M0, M1, and M2 conditions (24 h). **(E)** Immunofluorescence imaging of iNOS (green) and DAPI (blue) in WT and IIGP1-KO cells across M0, M1, and M2 conditions (24 h). scale bar = 10 μm. **(F)** Nitric oxide (NO) production (Griess assay) in WT and IIGP1-KO macrophages following 24 h stimulation with IFN-γ (100 ng/mL) or IL-4 (20 ng/mL). **(G)** Immunoblot of IIGP1 and iNOS in WT and IIGP1-KO cells treated with a dose-response range of IFN-γ (0-200 ng/mL, 24 h). (H) Griess assay measurement of NO production following the corresponding dose-response IFN-γ stimulation. **(I) Transmission electron microscopy (TEM)** images confirming the endpoint ultrastructure of WT and IIGP1-KO macrophages under M0 and M1 conditions (24 h); yellow arrows indicate the large intracellular granular vacuoles that dynamically emerge during the 16-18 h live-cell imaging window. Data are mean ± SD (n=3 independent biological replicates). Statistical significance was determined using an unpaired Student’s t-test (for panel E quantification) or one-way ANOVA (for all other panels) (p ≤ 0.05, ** p ≤ 0.01, *** p ≤ 0.001, **** p ≤ 0.0001; NS, not significant).

To ascertain whether this blunted iNOS induction reflects a downstream signaling threshold effect or an absolute cell-intrinsic requirement for IIGP1, WT and IIGP1-deficient macrophages were stimulated with graded concentrations of IFN-γ (0-200 ng/mL). In WT cells, IFN-γ induced a robust, dose-dependent increase in both IIGP1 and iNOS protein levels (**Fig. 3H**), alongside a parallel increase in nitrite accumulation (**Fig. 3I**). In contrast, IIGP1-deficient macrophages exhibited a total absence of IIGP1 across all doses, and iNOS induction remained severely blunted even at the highest concentration of IFN-γ tested (200 ng/mL) (**Fig. 3H**). Mirroring the protein expression data, NO production in the knockout cells remained impaired and was significantly lower than WT controls across the entire dose-response curve (**Fig. 3I**).

To capture the dynamics of this cellular remodeling, we performed 24-hour live-cell time-lapse imaging during M1 polarization. In WT macrophages, a pronounced accumulation of intracellular vacuoles emerged 16-18 hours post-IFN-γ stimulation **(Movie S1)**. In sharp contrast, this vacuolation process was severely blunted in IIGP1-KO cells **(Movie S2)**. Corroborating these real-time kinetics, endpoint transmission electron microscopy (TEM) at 24 hours confirmed abundant, distinct intracellular granular vacuoles in WT M1 macrophages (yellow arrows) that were markedly reduced in number and structural prominence in IIGP1-KO cells (**Fig. 3J**). Together, these live-imaging and ultrastructural data establish that IIGP1 is required for the dynamic vacuolar biogenesis characteristic of inflammatory M1 polarization.

These results demonstrate that the defect in iNOS expression and NO synthesis in IIGP1-deficient macrophages is not due to insufficient IFN-γ signaling, but rather reflects a cell-intrinsic requirement for IIGP1 downstream of the JAK-STAT1 pathway. Thus, these findings establish that IIGP1 is an essential cell-intrinsic regulator of the M1 macrophage inflammatory effector program.

### IIGP1 influences inflammatory macrophage architecture and polarization

Given that IIGP1 deficiency severely reduces iNOS induction and nitric oxide synthesis, we next investigated whether IIGP1 broadly orchestrates other facets of the inflammatory macrophage phenotype, including cellular architecture and transcriptional activation. Morphological assessment using rhodamine-phalloidin staining revealed that while wild-type (WT) M1 macrophages exhibit characteristic cellular spreading and an elongated, spindle-like morphology, IIGP1-deficient macrophages display markedly reduced spreading and altered cellular architecture following IFN-γ stimulation (**Fig. 4A**). Quantitative morphometric profiling confirmed these observations, revealing significantly reduced cell area, length, perimeter, and Feret diameters in IIGP1-deficient M1 macrophages compared to WT controls (**Fig. 4B**). Conversely, cellular solidity and circularity were significantly increased in the knockout cells, reflecting a rounder, unelongated morphology (**Fig. 4C**). At the transcriptional level, quantitative RT-PCR demonstrated that IIGP1-deficient M1 macrophages exhibit significantly reduced expression of key pro-inflammatory cytokines and chemokines, including Il6, Il1b, and Ccl2, relative to WT controls (**Fig. 4D**). Furthermore, flow cytometric analysis revealed a robust reduction in the surface expression of critical co-stimulatory and antigen-presentation markers, including MHC class II, CD86, and CD80, in the knockout cells (**Fig. 4E**). Collectively, these findings establish that IIGP1 is required to coordinate the broader M1 macrophage polarization program, governing cellular architecture, pro-inflammatory gene transcription, and surface marker upregulation.

**Figure 4.**
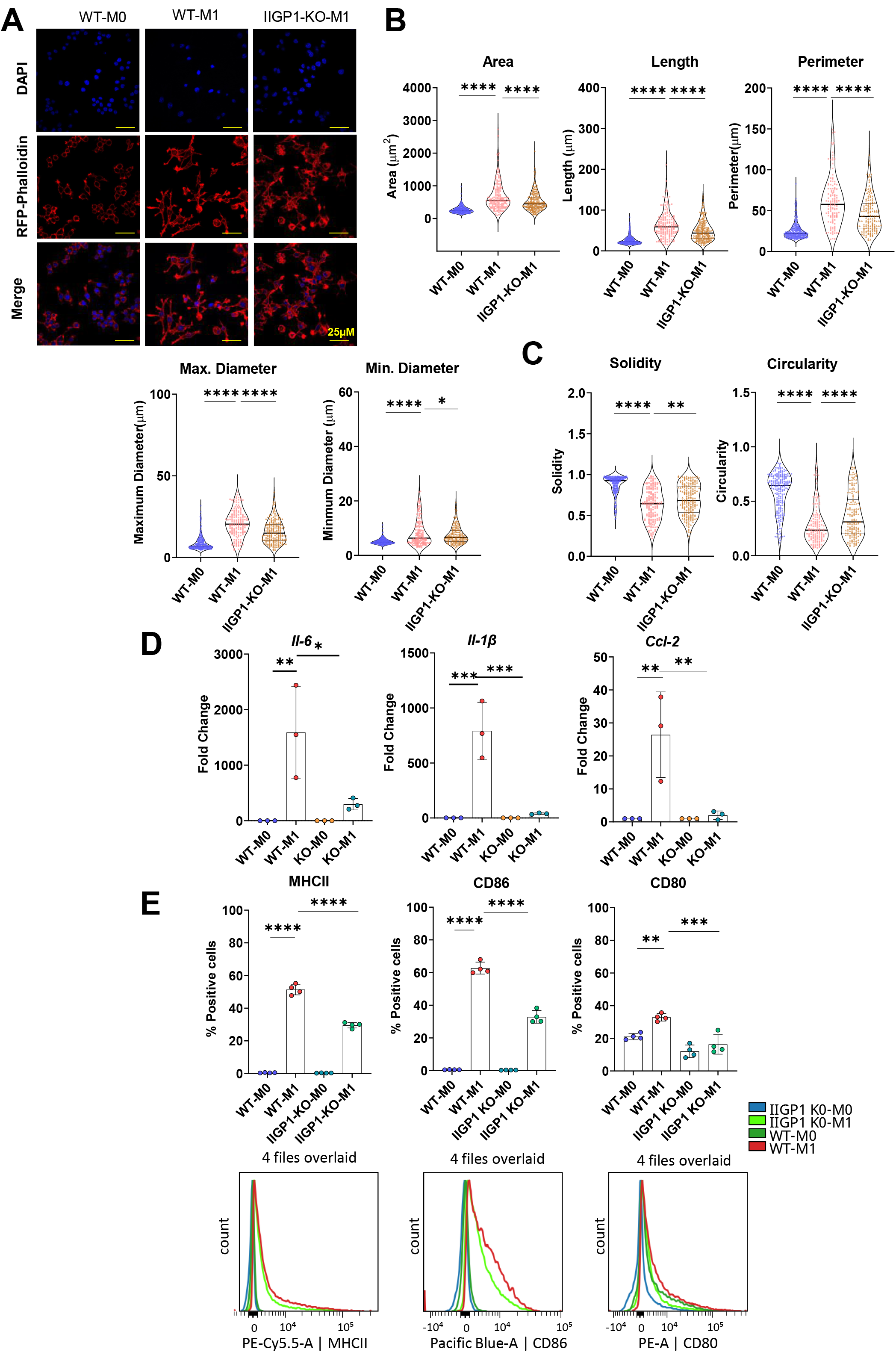
IIGP1 influences inflammatory macrophage architecture and activation. **(A)** Representative immunofluorescence images of WT and IIGP1-KO RAW264.7 macrophages under M0 and M1 (IFN-γ) conditions (24 h) stained with rhodamine-phalloidin (F-actin, red) and DAPI (nuclei, blue); scale bar = 25 μm. **(B)** Quantitative morphological profiling of WT-M1 and IIGP1-KO-M1 cells showing cell area, length, perimeter, and maximum/minimum Feret diameters (ImageJ analysis). **(C)** Derived morphological parameters quantifying solidity and circularity to assess cell shape and elongation. **(D)** Quantitative RT-PCR (qPCR) analysis of M1 inflammatory genes (*Il6*, *Il1b*, and *Ccl2*) in WT and IIGP1-KO macrophages across M0, M1, and M2 conditions, normalized to M0 controls. **(E)** Flow cytometric analysis of M1 surface markers (MHC class II, CD86, and CD80) in WT and IIGP1-KO cells under M0 and M1 conditions (24 h), displayed as percentage of positive cells (upper panels) and mean fluorescence intensity (MFI) (lower panels). Data represent mean ± SD (n=3 independent biological replicates). Statistical significance was determined by one-way ANOVA with Tukey’s post-hoc test (*p* ≤ is less than or equal to ≤ 0.05, ** *p* ≤ is less than or equal to ≤ 0.01, *** *p* ≤ is less than or equal to ≤ 0.001, **** *p* ≤ is less than or equal to ≤ 0.0001)

### IIGP1 associates with mitochondria and promotes metabolic remodeling in inflammatory macrophages

Since IIGP1 is induced downstream of IFN-γ-JAK-STAT1 signaling and is required for inflammatory macrophage polarization, we next sought to define the molecular mechanism by which it sustains the inflammatory phenotype. Because mitochondrial remodeling and metabolic reprogramming are fundamental to inflammatory macrophage polarization (19–20), we investigated whether IIGP1 functions through the regulation of mitochondrial pathways. To map the IIGP1 interactome, we performed immunoprecipitation coupled with mass spectrometry (IP-MS) in IFN-γ-stimulated RAW264.7 macrophages. Robust enrichment of IIGP1 was validated in the immunoprecipitates compared to IgG controls (**Fig. 5A**). Comparative proteomic analysis identified 557 proteins uniquely enriched in the IIGP1 interactome (**Fig. 5B**), and pathway enrichment analysis revealed a striking overrepresentation of mitochondrial and metabolism-associated proteins (**Fig. 5C**).

**Figure 5.**
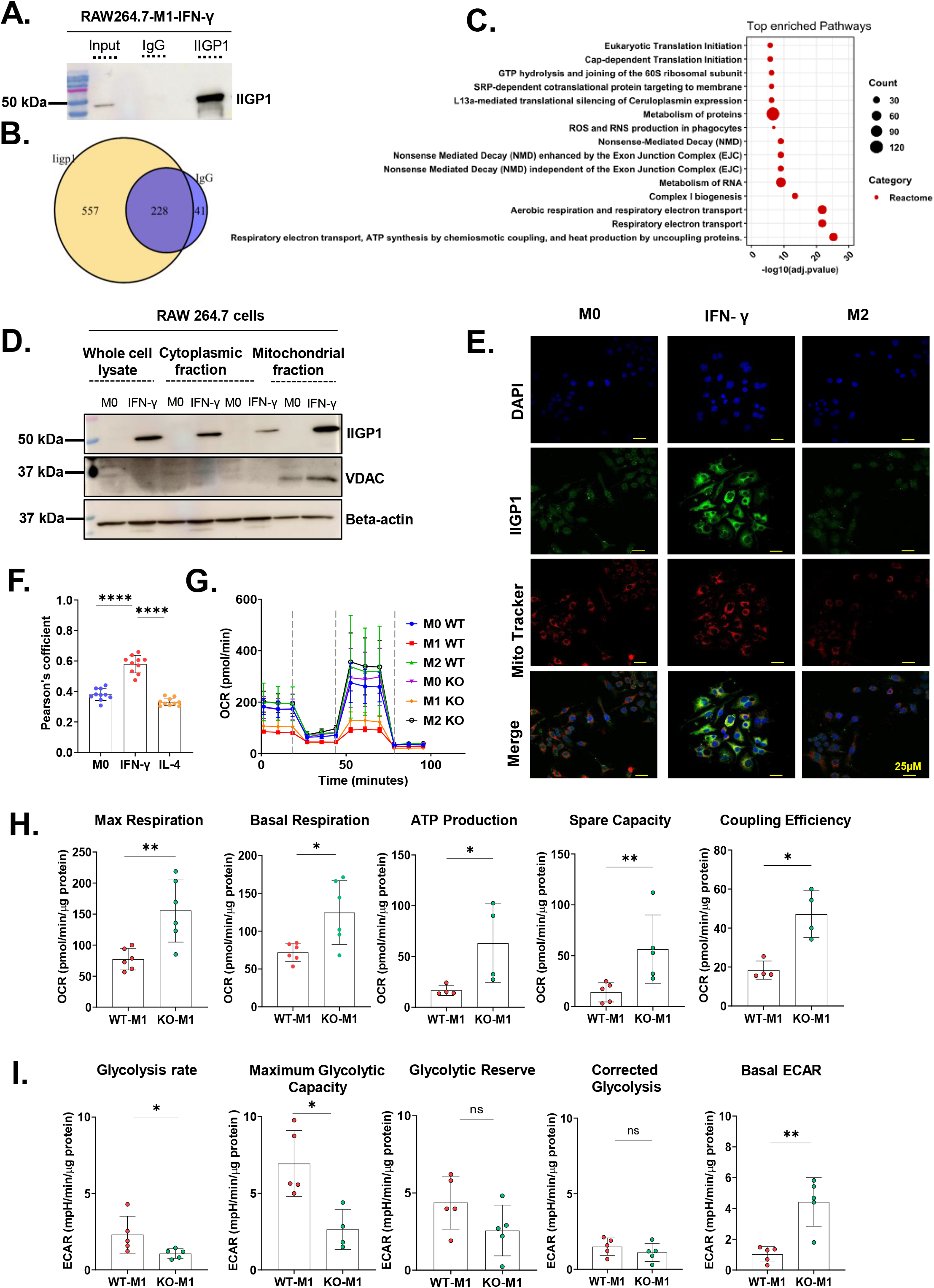
IIGP1 associates with mitochondria and promotes metabolic remodeling in inflammatory macrophages. **(A-C)** Mass spectrometry (MS) profiling of the IIGP1 interactome in RAW264.7 macrophages following IFN-γ stimulation (100 ng/mL, 24 h). **(A)** Immunoblot validation of IIGP1 immunoprecipitation (IP) across input, IgG control, and IP fractions. **(B)** Venn diagram illustrating unique and overlapping proteins identified between IgG control and IIGP1 pull-downs. **(C)** Pathway enrichment analysis of the 557 proteins uniquely associated with IIGP1. **(D)** Immunoblot of subcellular fractions from M0 and M1 macrophages probed for IIGP1, VDAC (mitochondrial marker), and β-actin (cytosolic/loading control). **(E)** Immunofluorescence imaging of IIGP1 (green), Mito Tracker Red (mitochondria), and DAPI (nuclei) in M1 and M2 polarized cells; scale bar = 25 μm. **(F)** Quantification of IIGP1-mitochondria colocalization. **(G, H)** Seahorse XF Mito Stress Test in WT and IIGP1-KO macrophages post-polarization (24 h) displaying **(G)** oxygen consumption rate (OCR) kinetic traces and **(H)** quantified respiratory parameters (basal, ATP-linked, spare respiratory capacity, coupling efficiency, and proton leak). **(I)** Seahorse XF Glycolysis Stress Test kinetic traces of extracellular acidification rate (ECAR) and quantified glycolytic parameters (glycolysis, glycolytic capacity, reserve, and basal ECAR). Data are mean ± SD (n=3-5 independent biological replicates). Statistical significance was determined using an unpaired Student’s t-test or one-way ANOVA (for all other panels) (p ≤ 0.05, ** p ≤ 0.01, *** p ≤ 0.001, **** p ≤ 0.0001; NS, not significant).

To experimentally validate this mitochondrial association, we first performed biochemical fractionation followed by immunoblotting of purified compartments, confirming the enrichment of IIGP1 within the mitochondrial fraction with minimal detection in the cytosolic compartment (**Fig. 5D**). Corroborating this, immunofluorescence imaging demonstrated pronounced colocalization of IIGP1 with Mito-tracker following IFN-γ stimulation (**Fig. 5E and F**), confirming its preferential mitochondrial localization during inflammatory polarization.

We next asked whether this mitochondrial association drives the metabolic remodeling characteristic of M1 macrophages. Initial screening of extracellular pH showed that while wild-type (WT) M1 macrophages exhibited expected acidification relative to M0 and M2 states, IIGP1-deficient M1 macrophages failed to undergo this shift **(Fig. S3A)**, suggesting impaired glycolytic reprogramming. To directly quantify cellular bioenergetics, we performed Seahorse extracellular flux analyses. Under M1-polarizing conditions, WT macrophages displayed a classic metabolic shift, characterized by suppressed oxidative phosphorylation (OXPHOS) and enhanced glycolytic activity. In sharp contrast, IIGP1-deficient M1 cells exhibited significantly elevated OXPHOS parameters, including increased basal respiration, ATP-linked respiration, and spare respiratory capacity (**Fig. 5G and H**). This sustained oxidative metabolism was accompanied by a reduction in glycolysis, glycolytic capacity, and glycolytic reserve (**Fig. 5I**). Collectively, these findings demonstrate that the loss of IIGP1 prevents the metabolic transition toward glycolysis and partially restores oxidative metabolism, indicating defective metabolic reprogramming. Together, these data identify IIGP1 as a critical mitochondria-associated effector downstream of IFN-γ signaling that directly orchestrates the bioenergetic shift required for inflammatory macrophage polarization.

### IIGP1 regulates mitochondrial redox state and mitochondrial organization during inflammatory macrophage polarization

Given the required role of IIGP1 in driving inflammatory macrophage polarization and metabolic reprogramming, we investigated its role in mitochondrial remodeling. Pro-inflammatory macrophage polarization typically repurposes mitochondria from ATP generation to inflammatory signaling via reactive oxygen species (mtROS) production, membrane potential (Ψ_m_) alterations, and structural dynamics (19–20).

To evaluate mitochondrial mass, we performed Mitotracker staining. While wild-type (WT) M1 macrophages showed increased mass relative to M0 controls, IIGP1-KO-M1 macrophages exhibited a paradoxical decrease in mass remaining comparable to M0 cells. This indicates impaired mitochondrial biogenesis during inflammatory activation (**Fig. 6A**). DCFDA and MitoSOX staining revealed that WT inflammatory macrophages exhibited robust increases in total cellular and mitochondrial ROS, whereas IIGP1 deficiency significantly blunted these signals (**Fig. 6B**). This establishes that IIGP1 is indispensable for the mitochondrial redox remodeling required to drive pro-inflammatory gene networks.

**Figure 6.**
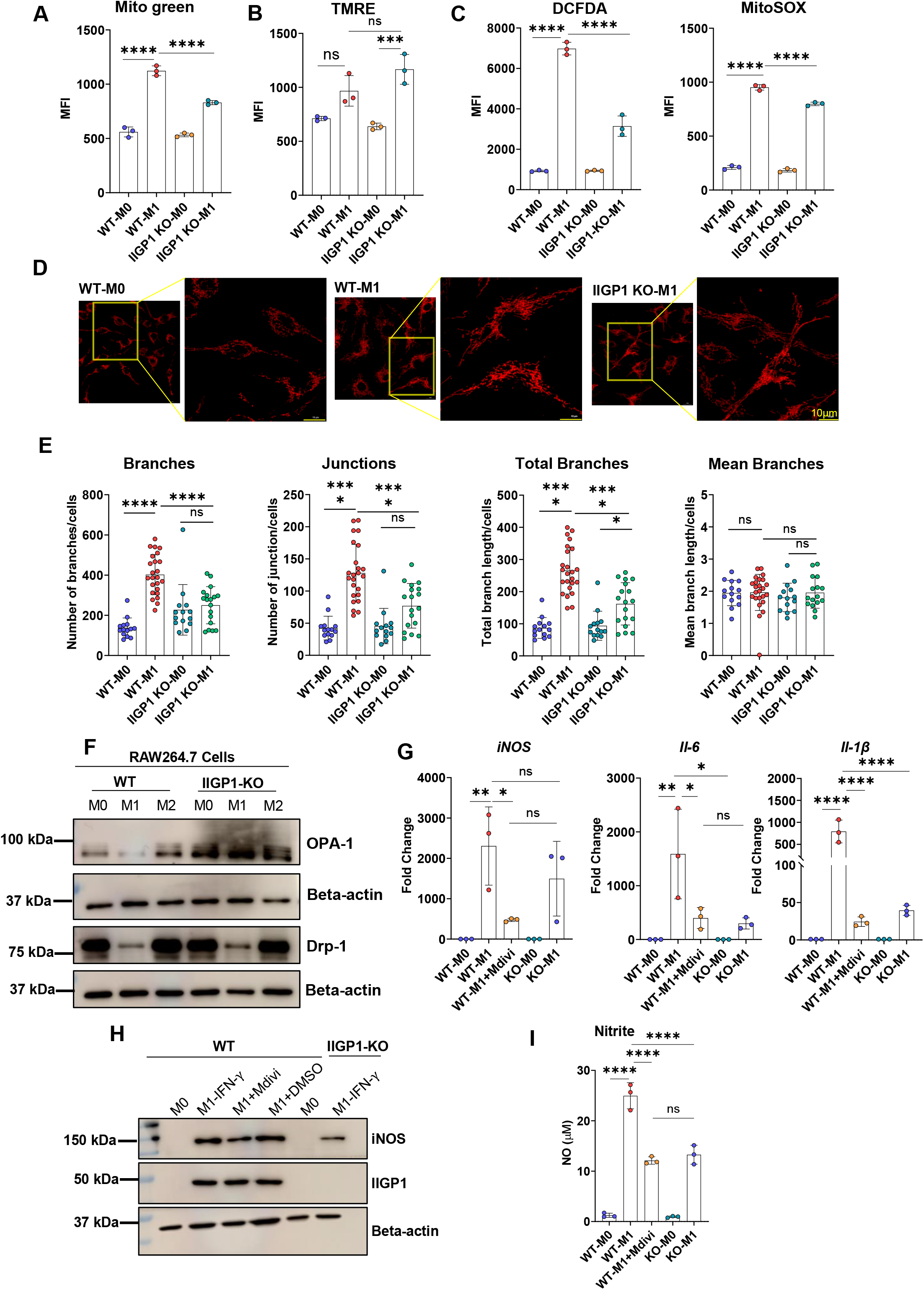
IIGP1 regulates mitochondrial redox state and structural organization during inflammatory macrophage activation. **(A-C)** Flow cytometric profiling of mitochondrial physiology in WT and IIGP1-KO RAW264.7 macrophages under M0 and M1 (IFN-γ) polarizing conditions (24 h). Panels show representative mean fluorescence intensity (MFI) quantification for **(A)** mitochondrial mass (Mito Tracker-Green), **(B)** intracellular ROS (DCFDA) and mitochondrial superoxide (MitoSOX^™^), and **(C)** mitochondrial membrane potential (TMRE). **(D)** Confocal immunofluorescence imaging of mitochondrial morphology (Mito Tracker^™^ Red) in WT and IIGP1-KO cells during M0 and M1 polarization. **(E)** Quantitative morphometric analysis of mitochondrial networks skeletonized from confocal images using ImageJ. **(F)** Immunoblot analysis of mitochondrial fission (DRP1) and fusion (OPA1) proteins across M0, M1, and M2 polarizing states; β-actin served as the loading control. **(G, H)** Functional evaluation of WT and IIGP1-KO cells pretreated with Mdivi-1 (25 μM) prior to 24-h IFN-γ stimulation, showing **(G)** quantitative RT-PCR (qPCR) analysis of M1-associated transcripts (*Nos2*, *Il6*, and *Il1b*), and **(H)** iNOS and IIGP1 immunoblots accompanied by quantification of culture supernatant nitric oxide (NO) production via Griess assay. Data are mean ± SD (n=3 independent biological replicates). Statistical significance was determined using a one-way ANOVA (for all other panels) (p ≤ 0.05, ** p ≤ 0.01, *** p ≤ 0.001, **** p ≤ 0.0001; NS, not significant).

To evaluate bioenergetic status, we measured Ψ_m_ via TMRE staining. IIGP1-KO-M1 macrophages demonstrated significantly higher TMRE fluorescence than WT controls (**Fig. 6C**). This hyperpolarization aligns with the enhanced oxidative phosphorylation (OXPHOS) and impaired glycolysis previously observed in these cells (**Fig. 5G, H and I)**, confirming that IIGP1 loss shifts mitochondria toward a hyper-energized, oxidative state. We next examined mitochondrial morphology using fluorescence microscopy (**Fig. 6D**). Inflammatory WT-M1 macrophages displayed an interconnected network with increased branching and junctions compared to WT-M0 cells. In contrast, IIGP1-KO-M1 macrophages exhibited a simplified architecture with fewer branches and junctions resembling WT-M0 cells, mirroring their reduced cellular spreading (**Fig. 4A**). Quantitative morphometry confirmed this shift, showing a significant reduction in network branches and junctions in IIGP1-K0-M1 macrophages (**Fig. 6E**). To determine the molecular basis underlying these structural changes, we examined the expression of the fusion protein OPA1 and the fission protein DRP1. Immunoblot analysis revealed increased OPA1 expression levels in IIGP1-deficient macrophages (**Fig. 6F**), suggesting a molecular shift toward enhanced fusion in the absence of IIGP1 that supports a fused mitochondrial state and suppresses the inflammatory macrophage program.

To determine whether these altered dynamics contribute to the inflammatory defects observed following IIGP1 loss, we pharmacologically inhibited mitochondrial fission using the DRP1 inhibitor Mdivi-1. We reasoned that blocking DRP1-mediated fission, a hallmark remodeling event in M1 macrophages, would shift the structural balance toward OPA1-driven fusion, thereby altering macrophage inflammatory activation. Underscoring this mechanism, Mdivi-1 treatment in WT M1 macrophages significantly attenuated the transcript expression of *Nos2*, *Il-6*, and *Il-1β* (**Fig. 6G**). In line with these transcriptional defects, Mdivi-1 treatment also suppressed iNOS protein expression and subsequent nitric oxide (NO) production, effectively phenocopying the functional impairments observed in IIGP1-deficient macrophages (**Fig. 6H and I**).

Collectively, these findings demonstrate that IIGP1 is required for coordinated mitochondrial remodeling during inflammatory macrophage activation. Loss of IIGP1 disrupts mitochondrial metabolic adaptation, hyperpolarizes membrane potential, impairs redox homeostasis, and shifts mitochondrial dynamics toward a fused, oxidative state, consequently compromising the inflammatory macrophage program. These findings place mitochondrial remodeling downstream of IFN-γ-IIGP1 signaling as a key mechanism supporting inflammatory macrophage effector function.

### IIGP1 is required for the antibacterial effector function of inflammatory macrophages

Having established that IIGP1 is required for inflammatory macrophage activation and mitochondrial remodeling, we next investigated whether these cellular defects compromise the primary antibacterial effector functions of the cell. Specifically, we evaluated whether IIGP1 expression is mandatory for the clearance and destruction of intracellular pathogens. Using intracellular colony-forming unit (CFU) assays, we first measured the clearance of *Escherichia coli* DH5α. We observed a significantly higher recovery of viable intracellular bacteria from IIGP1-deficient macrophages compared with wild-type (WT) controls at 2 hours post-infection (**Fig. 7A**), demonstrating an impairment in early bacterial clearance following IIGP1 loss. To determine if this defect extended to virulent intracellular pathogens, we tracked the survival of *Salmonella* Typhimurium. The intracellular killing capacity was similarly compromised in IIGP1-deficient macrophages, as evidenced by a significantly higher intracellular bacterial burden at both the 2-hour (**Fig. 7B**) and 7-hour (**Fig. 7C**) post-infection time points. This persistent failure to restrict bacterial replication aligns with the defective inflammatory phenotype observed during the activation phase of these knockout cells. Based on our results, we propose a model for IIGP1-dependent immunometabolic reprogramming during macrophage activation (**Fig. 7D**). In WT macrophages, IFN-γ-driven IIGP1 expression suppresses OPA1, facilitating the mitochondrial structural fragmentation, reactive oxygen species (mtROS) production, and glycolytic flux required for robust M1 pro-inflammatory polarization and intracellular bacterial clearance. Conversely, IIGP1 deficiency increases OPA1 expression, shifting mitochondria toward a hyperpolarized (ΔΨ*m*), highly fused oxidative state. This structural and metabolic disruption severely blunts pro-inflammatory gene induction (iNOS, cytokines, costimulatory markers) and compromises the host antibacterial response.

**Figure 7.**
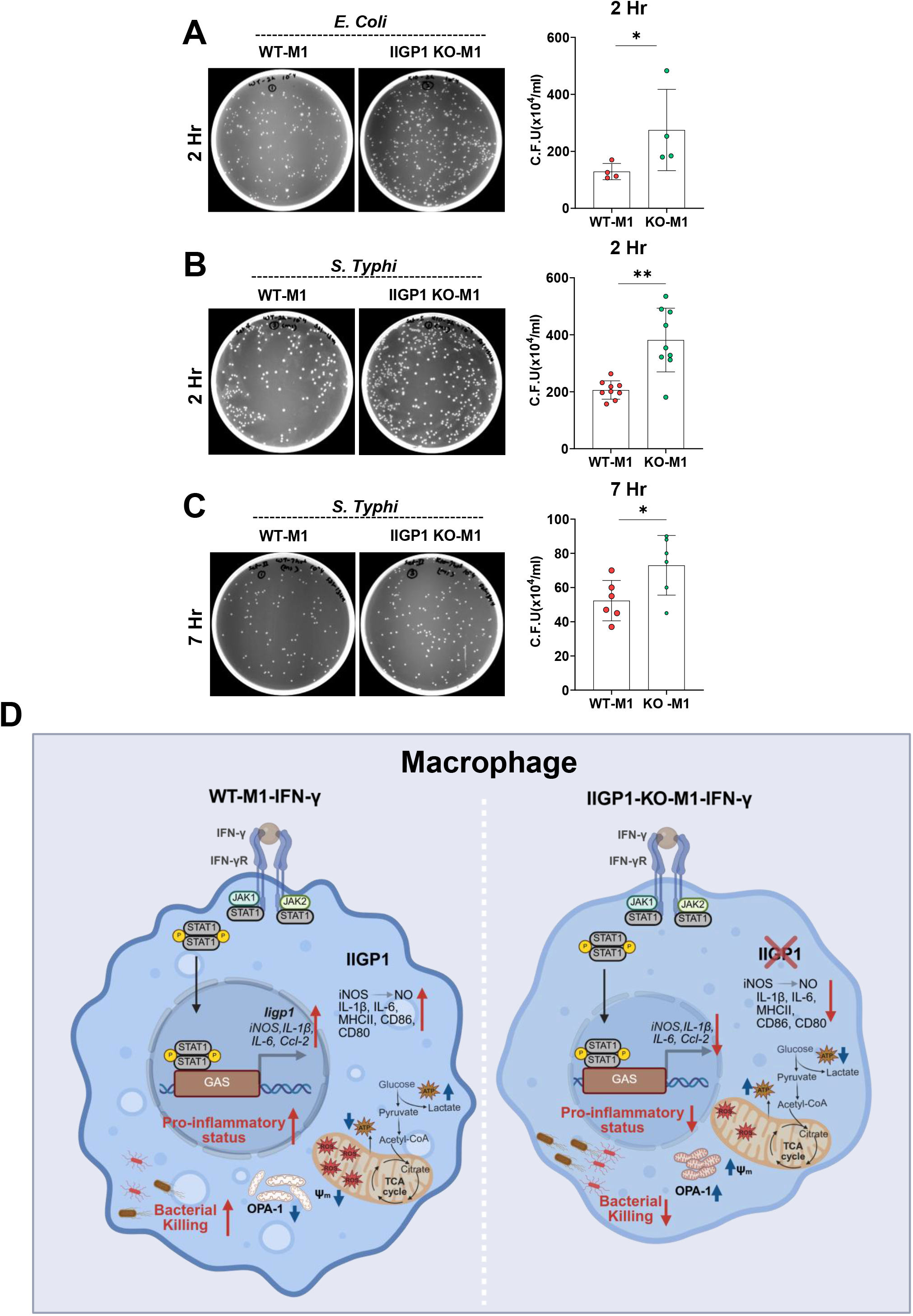
IIGP1 is required for the antibacterial effector function of inflammatory macrophages. **(A-C)** RAW264.7 wild-type (WT) and IIGP1-knockout (KO) macrophages were polarized to the M1 phenotype with IFN-γ (24 h) and infected with *Escherichia coli* DH5α or *Salmonella* Typhi (MOI 1:40) for 1 h. Extracellular bacteria were eliminated via gentamicin protection assay, and surviving intracellular bacteria were determined by colony-forming unit (CFU) agar plating. **(A)** Representative LB agar plates and corresponding CFU quantification of intracellular *E. coli* DH5α recovered 2 h post-infection. **(B)** Representative agar plates and CFU quantification of intracellular *S.* Typhi recovered 2 h post-infection. **(C)** Representative agar plates and CFU quantification of intracellular *S.* Typhi recovered 7 h post-infection. **(D)** Schematic model of IIGP1-dependent immunometabolic reprogramming during macrophage activation. In WT cells (left), IFN-γ-induced IIGP1 suppresses OPA1, facilitating the mitochondrial structural fragmentation, reactive oxygen species (mtROS) production, and glycolytic flux required for robust M1 pro-inflammatory polarization and bacterial clearance. Conversely, IIGP1 deficiency (right) increases OPA1 expression, shifting the network toward a hyperpolarized (ΔΨ*m*), highly fused oxidative state that blunts pro-inflammatory mediator expression and compromises host defense. Data are presented as mean ± SD from three independent experiments. Statistical analysis was performed using the Mann-Whitney test (\**p* = 0.029; \*\**p* = 0.0079; NS, not significant).

## Discussion

Inflammatory macrophage polarization requires the coordinated integration of cytokine signaling, mitochondrial remodeling, and metabolic reprogramming; however, the molecular effectors that couple IFN-γ signaling to these mitochondrial adaptations have remained poorly defined. In this study, we identify IIGP1 as an IFN-γ-inducible molecular effector that connects JAK-STAT1 inflammatory signaling to mitochondrial homeostasis and immunometabolic reprogramming in macrophages. Using quantitative proteomics, we identified IIGP1 as one of the most robustly induced proteins during inflammatory macrophage polarization. Functional characterization demonstrated that IIGP1 operates downstream of IFN-γ-JAK-STAT1 signaling and is required for the establishment of the inflammatory macrophage phenotype, including iNOS induction, nitric oxide production, pro-inflammatory gene expression, cytokine secretion, and macrophage architectural remodeling. IIGP1 associates with mitochondria and supports the glycolytic metabolic reprogramming, redox adaptation, membrane potential regulation, and structural remodeling characteristic of inflammatory macrophages. Consistent with these mitochondrial and metabolic defects, loss of IIGP1 compromised effective intracellular bacterial killing. Collectively, these findings address a previously unrecognized gap in our understanding of how IFN-γ signaling is translated into organelle-level remodeling and metabolic adaptation in inflammatory macrophages.

Our data establish that IIGP1 expression during inflammatory macrophage activation is governed by the IFN-γ-JAK-STAT1 signaling axis. STRING protein interaction network analysis identified a functional association between IIGP1 and STAT1, and in silico promoter analysis revealed putative STAT1 binding motifs within the IIGP1 promoter region, suggesting possible transcriptional regulation by STAT1. Pharmacological inhibition of JAK signaling using tofacitinib markedly attenuated IIGP1 induction at both transcript and protein levels, confirming that IIGP1 is a JAK-STAT1-dependent IFN-γ target gene. These findings are consistent with previous studies demonstrating that members of the IRG family are among the most strongly and rapidly induced IFN-γ target genes in macrophages (27–35). The IFN-γ-JAK-STAT1 axis is known to induce a broad transcriptional program that promotes antimicrobial and inflammatory macrophage functions (25); however, the identities of downstream effector proteins that translate this transcriptional response into organelle-level remodeling have remained incompletely characterized (23–24). The identification of IIGP1 as a STAT1-dependent gene that regulates mitochondrial homeostasis, therefore, provides a previously unrecognized link between the IFN-γ transcriptional response and mitochondrial metabolic adaptation in inflammatory macrophages. Future studies employing chromatin immunoprecipitation assays will be important to confirm direct STAT1 occupancy at the IIGP1 promoter and define the precise regulatory elements governing its IFN-γ-dependent induction.

IIGP1-deficient macrophages exhibited broad impairment of the inflammatory macrophage program, including significantly reduced iNOS expression, nitric oxide production, pro-inflammatory cytokine and chemokine expression, and surface expression of co-stimulatory and antigen presentation markers. The impairment of iNOS induction and nitric oxide production is particularly significant, given the central role of the iNOS-NO axis in macrophage antimicrobial defense, inflammatory amplification, and pathogen clearance (1–4). The dose-response experiment demonstrated that escalating IFN-γ concentrations could not rescue iNOS induction in IIGP1-deficient macrophages, firmly establishing that the defect reflects a cell-intrinsic requirement for IIGP1 downstream of cytokine receptor activation rather than a signaling threshold effect. This places IIGP1 functionally downstream of JAK-STAT1 activation and upstream of iNOS transcriptional induction, suggesting that IIGP1 may act as a signal transduction intermediate that connects IFN-γ-dependent transcriptional output to the execution of the M1 effector program. IIGP1-deficient macrophages exhibited reciprocally elevated ARG1 expression and urea production under alternative activation conditions, suggesting that IIGP1 deficiency shifts the polarization balance toward an M2-like state. This reciprocal shift is consistent with the well-established metabolic competition between iNOS and ARG1 for their shared substrate L-arginine (10) and raises the possibility that IIGP1 contributes to maintaining the metabolic and transcriptional asymmetry between M1 and M2 polarization states. Furthermore, the morphological defects observed in IIGP1-deficient macrophages-including reduced cellular spreading, altered architecture, and rounder cellular morphology-are consistent with previous reports linking cytoskeletal and morphological remodeling to inflammatory macrophage polarization and effector function (44–46).

A central finding of this study is the mitochondrial association of IIGP1 during inflammatory macrophage activation. IP-MS analysis identified significant enrichment of mitochondrial and metabolism-associated proteins among IIGP1-interacting partners, and both immunofluorescence colocalization and biochemical fractionation independently confirmed mitochondrial enrichment of IIGP1 following IFN-γ stimulation. These observations extend the previously characterized biology of IIGP1, which has been reported to localize predominantly to the endoplasmic reticulum and Golgi apparatus, where it participates in intracellular membrane organization and antimicrobial responses. The present findings raise the possibility that IIGP1 undergoes dynamic redistribution to mitochondria specifically during inflammatory activation, consistent with its role as an IFN-γ-responsive effector that participates in membrane remodeling across multiple intracellular compartments. The human IRG family member IRGM has been shown to associate with mitochondria and regulate mitochondrial dynamics, mitochondrial quality control, and autophagy-related processes (36–37), suggesting that mitochondrial association may represent a conserved functional feature shared among IRG family members. The mitochondrial localization of IIGP1 is also consistent with its established role in intracellular membrane remodeling (36–37) and suggests that IIGP1 may contribute to the remodeling of mitochondrial membranes that accompanies inflammatory macrophage polarization. The identity of specific mitochondrial interaction partners of IIGP1 revealed by IP-MS analysis represents an important focus for future mechanistic investigation aimed at defining the precise molecular mechanism by which IIGP1 regulates mitochondrial homeostasis.

One of the most striking consequences of IIGP1 deficiency was the failure of macrophages to undergo the glycolytic metabolic shift that characterizes inflammatory macrophage activation. Wild-type M1 macrophages exhibited the expected reduction in oxidative phosphorylation parameters and elevation in glycolytic activity, consistent with the well-established Warburg-like metabolic reprogramming of inflammatory macrophages (12–13). In contrast, IIGP1-deficient M1 macrophages displayed significantly elevated mitochondrial respiratory activity and impaired glycolytic adaptation, indicating a fundamental failure to execute the metabolic switch required for inflammatory activation. This metabolic defect was further corroborated by extracellular pH measurements, which demonstrated that IIGP1-deficient macrophages failed to exhibit the increased acidification characteristic of glycolytically active M1 macrophages. These findings are consistent with growing evidence demonstrating that metabolic reprogramming is not merely a downstream consequence of inflammatory activation but is functionally required for the full execution of the inflammatory macrophage effector program (18–20). The impaired glycolytic reprogramming in IIGP1-deficient macrophages likely contributes mechanistically to the reduced iNOS expression and nitric oxide production observed in these cells, given the established dependence of sustained iNOS activity on glycolytic metabolic support and the availability of glycolytic intermediates as precursors for NO biosynthesis (47–48). Together, these data position IIGP1 as a previously unrecognized regulator of the immunometabolic switch in inflammatory macrophages, operating at the interface of IFN-γ signaling and mitochondrial metabolic adaptation.

Beyond metabolic reprogramming, IIGP1 deficiency broadly disrupted mitochondrial homeostasis in inflammatory macrophages, affecting mitochondrial content, redox state, membrane potential, structural organization, and dynamics regulatory proteins. IIGP1-deficient M1 macrophages exhibited significantly reduced mitochondrial content, as evidenced by decreased Mito Tracker Green fluorescence, suggesting impaired mitochondrial mass during inflammatory activation. Additionally, mitochondrial and total cellular ROS levels were significantly reduced in IIGP1-deficient macrophages, consistent with their failure to undergo glycolytic reprogramming and their retention of oxidative metabolic characteristics. Mitochondrial ROS have been shown to play important signaling roles in inflammatory macrophage activation, including amplification of NF-κB-dependent inflammatory gene expression and support of antimicrobial effector mechanisms (20–22). The reduced ROS levels in IIGP1-deficient macrophages may therefore contribute both to the impaired inflammatory gene expression and to the defective antibacterial killing capacity observed in these cells. Elevated mitochondrial membrane potential in IIGP1-deficient macrophages further supported a shift toward oxidative metabolism, consistent with Seahorse flux analysis data demonstrating elevated oxidative phosphorylation in these cells. At the structural level, IIGP1-deficient macrophages exhibited a compact, less branched mitochondrial network accompanied by altered expression of the dynamic regulatory proteins OPA1 and DRP1, suggesting dysregulation of the mitochondrial fusion-fission balance in the absence of IIGP1. Pharmacological inhibition of DRP1-mediated fission using Mdivi-1 phenocopied the inflammatory defects observed in IIGP1-deficient macrophages-reducing iNOS expression, nitric oxide production, and pro-inflammatory gene expression in wild-type M1 macrophages - while failing to further suppress these parameters in IIGP1-deficient cells. This epistatic relationship supports the conclusion that IIGP1 and DRP1-mediated mitochondrial fission operate within the same regulatory axis, and that IIGP1-dependent regulation of mitochondrial dynamics is functionally required for the establishment of the inflammatory macrophage program. These findings are consistent with emerging evidence demonstrating that mitochondrial dynamics are actively regulated during macrophage activation and contribute to inflammatory effector function (14–16).

The functional importance of IIGP1-dependent mitochondrial programming was directly demonstrated by the impaired antibacterial killing capacity of IIGP1-deficient macrophages. These cells exhibited significantly impaired intracellular killing of both *Escherichia coli* and *Salmonella Typhimurium* at early and late time points, demonstrating that the antibacterial defect is specific to killing mechanisms rather than phagocytic capacity. The antibacterial killing defect observed in IIGP1-deficient macrophages is likely attributable, at least in part, to the impaired iNOS induction and nitric oxide production observed in these cells, given the well-established and critical role of NO in macrophage bactericidal activity against both Gram-negative and Gram-positive pathogens (1–4). Reduced mitochondrial ROS generation may additionally contribute to the impaired bacterial killing, consistent with the established role of mitochondrial ROS as direct antimicrobial effectors and as amplifiers of inflammatory signaling in macrophages (20–22). These findings are consistent with the established role of IRG family members as important mediators of cell-autonomous host defense against a broad range of intracellular pathogens (27–35) and identify IIGP1 as a functionally important antimicrobial effector whose activity depends on mitochondrial metabolic programming rather than direct pathogen recognition or phagocytic function. Together, these observations establish a functional axis linking IFN-γ signaling through IIGP1 to mitochondrial remodeling and ultimately to effective macrophage antibacterial immunity.

Several important questions remain to be addressed in future studies. First, while our findings establish that IIGP1 associates with mitochondria and is required for mitochondrial remodeling, the precise molecular mechanism by which IIGP1 regulates mitochondrial dynamics and metabolic reprogramming remains to be defined. Whether IIGP1 directly interacts with components of the mitochondrial fission or fusion machinery, such as DRP1 or OPA1, or acts indirectly through regulation of mitochondrial membrane organization and lipid composition, will require detailed structural and biochemical investigation. Second, the identity of the specific mitochondrial binding partners of IIGP1, revealed by IP-MS analysis, warrants further investigation to define the molecular basis of its mitochondrial recruitment and function. Third, while the present studies were conducted primarily in RAW264.7 macrophages and supported by bone marrow-derived macrophage data, validation of key findings in primary macrophage systems and in vivo infection models will be important for establishing the full physiological relevance of IIGP1-dependent mitochondrial programming. Fourth, the human relevance of these findings is an important consideration. IIGP1 is a murine-specific p47 IRG family member, and whether the human IRG family member IRGM, which shares functional properties including mitochondrial association and regulation of autophagy (36,43) and performs analogous roles in human macrophage immunometabolism, warrants investigation. Finally, whether IIGP1-dependent mitochondrial programming contributes to macrophage function in inflammatory disease contexts beyond infection – including autoimmune diseases, metabolic inflammation, or conditions characterized by dysregulated macrophage activation – represents an important avenue for future exploration.

In summary, the present study identifies IIGP1 as an IFN-γ-inducible molecular effector that connects JAK-STAT1 inflammatory signaling to mitochondrial homeostasis and immunometabolic reprogramming in macrophages. Our findings demonstrate that IIGP1 is required for the coordinated mitochondrial remodeling, glycolytic metabolic adaptation, and effector programming that define the inflammatory macrophage phenotype, and that IIGP1-dependent mitochondrial programming is functionally required for effective intracellular bacterial killing. These findings advance our understanding of how IFN-γ signaling is translated into organelle-level remodeling and metabolic reprogramming and identify the IRG GTPase family as an important and previously unrecognized node connecting inflammatory cytokine signaling to mitochondrial immunometabolism. More broadly, these findings contribute to the emerging concept that mitochondrial remodeling is not merely a metabolic adaptation but a functionally integrated and actively regulated component of the inflammatory macrophage polarization program, with direct consequences for antimicrobial host defense.

## Experimental Procedures

### Cell culture and macrophage polarization

#### Generation of Bone Marrow-Derived Macrophages (BMDMs)

Bone marrow-derived macrophages (BMDMs) were generated from six- to eight-week-old male C57BL/6J mice obtained from the Experimental Animal Facility, National Centre for Cell Science (NCCS), Pune, India. All animal experiments were performed in accordance with the guidelines approved by the Institutional Animal Ethics Committee (IAEC) of NCCS, Pune. Bone marrow cells were isolated by flushing femurs and tibiae with sterile phosphate-buffered saline (PBS). Mononuclear cells were enriched by density gradient centrifugation using Histopaque^®^ (Cat. #1077, Sigma-Aldrich, USA) and resuspended in α-minimum essential medium (α-MEM; Gibco) supplemented with 10% fetal bovine serum (Cat. #16000-044, Gibco, USA) and 7 ng/mL macrophage colony-stimulating factor (M-CSF). Cells were cultured overnight in T-25 tissue culture flasks to allow adherence of stromal cells. The non-adherent bone marrow precursor cells were subsequently collected and differentiated into macrophages by culturing in α-MEM containing 10% FBS supplemented with 40 ng/mL M-CSF for 5 days. Fully differentiated unstimulated macrophages (M0) were used for subsequent polarization experiments.

#### RAW264.7 Cell Culture

Murine RAW264.7 macrophage-like cells (ATCC TIB-71) were obtained from the American Type Culture Collection (ATCC, Manassas, VA, USA). Cells were maintained in Dulbecco’s Modified Eagle Medium (DMEM) (Cat. #12100-046, Gibco, USA) supplemented with 10% (v/v) heat-inactivated fetal bovine serum (FBS) (Cat. #16000-044, Gibco, USA) and 1% penicillin-streptomycin (Penicillin-Streptomycin, 10,000 U/mL Cat. #15140122, Gibco, USA). Cells were cultured at 37°C in a humidified atmosphere containing 5% CO₂ and passaged every 2-3 days to maintain exponential growth.

#### Macrophage Polarization

Bone marrow-derived macrophages (BMDMs) and RAW264.7 macrophages were cultured under standard conditions prior to polarization. For inflammatory macrophage activation, cells were stimulated with recombinant murine IFN-γ (100 ng/mL; Cat. #315-05, PeproTech^®^, USA) for 24 h to generate M1 (IFN-γ) macrophages. Where indicated, macrophages were stimulated with lipopolysaccharide (lipopolysaccharides from Escherichia coli O111:B4; 10 ng/mL; Cat. # L2630, Sigma-Aldrich, USA) for 24 h to generate M1(LPS) macrophages. Alternatively, activated macrophages were generated by treatment with recombinant murine IL-4 (10 ng/mL; Cat. # 214-14, PeproTech^®^, USA) for 24 h to obtain M2 (IL-4) macrophages. Unstimulated macrophages cultured under identical conditions served as M0 controls. All cells were maintained at 37°C in a humidified incubator containing 5% CO₂.

### RNA Isolation and Quantitative Real-Time PCR (qRT-PCR)

Total RNA was isolated from unstimulated (M0) and polarized macrophages using TRIzol reagent (Cat. #15596018, Invitrogen, USA) according to the manufacturer’s instructions. RNA concentration and purity were determined spectrophotometrically. First-strand cDNA was synthesized from 1.2 μg of total RNA using a High-Capacity cDNA Reverse Transcription Kit (Cat. #4368814, Applied Biosystems, USA) following the manufacturer’s protocol. Quantitative real-time PCR (qRT-PCR) was performed using Power SYBR™ Green PCR Master Mix (Cat. #A25742, Applied Biosystems, USA) on a CFX96^™^ Real-Time PCR System (Bio-Rad). The cycling conditions consisted of an initial incubation at 50°C for 2 min and 95°C for 10 min, followed by 40 cycles of 95°C for 15 sec, 65°C for 30 sec, and 72°C for 30 sec. Relative gene expression was calculated using the 2^−ΔΔCt^ method with Actb (β-actin) as the internal reference gene. Gene expression levels were normalized to β-actin and expressed relative to unstimulated (M0) macrophages. Primer sequences are listed in Table 1.

### Western Blot

Cells were lysed on ice for 30 min in lysis buffer containing 20 mM Tris-HCl (pH 7.4), 150 mM NaCl, 1% Triton X-100, 10% glycerol, 0.1% SDS, 1 mM EDTA, 1 mM sodium orthovanadate, and 25 mM β-glycerophosphate supplemented with 1 mM phenylmethylsulfonyl fluoride (PMSF) and (Cat. #329986, Affymetrix, USA) protease inhibitor cocktail (Cat. #0403877013008, Merk, USA). Lysates were clarified by centrifugation at 11,500 × g for 20 min at 4°C, and protein concentrations were determined using the Bio-Red Protein Assay Dye Reagent (Cat. #5000006, Bio-Rad, USA). Equal amounts of protein (20-30 μg) were mixed with sample buffer, boiled at 95°C for 10 min, separated by 10% SDS-PAGE, and transferred onto PVDF membrane (Cat. #IPCH00010, Millipore, USA). Membranes were blocked with 5% (w/v) non-fat dry milk in TBST for 1 h at room temperature and incubated overnight at 4°C with the indicated primary antibodies (Table 2) diluted in 5% (w/v) BSA (Cat. #160069, MP-Biomedical, USA). After washing three times with TBST, membranes were incubated with HRP-conjugated goat anti-rabbit or goat anti-mouse secondary antibodies (Table 2) for 1 h at room temperature. Protein bands were detected using an enhanced chemiluminescence (ECL) substrate (Cat. #170-5061, #170-5062, Bio-Rad, USA) and visualized using an Amersham^™^ Image Quant^™^ 800 (Cytiva). Band intensities were quantified using ImageJ software (NIH), normalized to the corresponding loading control, and expressed relative to the control group within each experiment.

### CRISPR/Cas9-Mediated Generation of IIGP1-Deficient RAW264.7 Macrophages

IIGP1-deficient RAW264.7 macrophages were generated using CRISPR/Cas9-mediated genome editing. A single-guide RNA (sgRNA) targeting exon 2 of the murine Iigp1 (Irga6) gene was designed using the Genetic Perturbation Platform (Broad Institute) based on predicted on-target activity and off-target potential. The sgRNA sequence was cloned into the pCRISPR-CG01 vector at the BamHI and XbaI restriction sites. RAW264.7 cells were transfected with the resulting pCRISPR-CG01-sgIigp1 construct using Lipofectamine 3000 (Invitrogen, Cat. #L3000-001) according to the manufacturer’s instructions. Twenty-four hours after transfection, cells were selected with puromycin (Sigma-Aldrich, Cat. #P7255) (2 μg/mL) for 48 h. Surviving cells were subsequently expanded and subjected to limiting dilution to generate single-cell-derived clones. Individual clones were screened for IIGP1 expression following IFN-γ stimulation (100 ng/mL, 24 h). Loss of IIGP1 expression was initially assessed by immunofluorescence microscopy and subsequently confirmed by immunoblotting using an anti-IIGP1 antibody. Clones exhibiting complete loss of IIGP1 protein expression following IFN-γ stimulation were selected for subsequent experiments. The sgRNA sequence used for Iigp1 targeting is provided in Table 3.

### Nitric Oxide (NO) Production Assay

Nitric oxide (NO) production was quantified by measuring nitrite accumulation in culture supernatants using the Griess assay. RAW264.7 macrophages were seeded in 96-well plates at 1.5 × 10⁵ cells per well and allowed to adhere overnight. Cells were either left unstimulated (M0) or polarized under the indicated experimental conditions, including stimulation with 100 ng/mL IFN-γ for 24 h, unless otherwise stated. Following stimulation, culture supernatants were collected and centrifuged briefly to remove cellular debris. Equal volumes (50 μL) of culture supernatant and Griess reagent (Cat. #G4410, Sigma-Aldrich) were mixed in a fresh 96-well plate and incubated for 10 min at room temperature protected from light. Absorbance was measured at 540 nm using an Infinite^®^ F200 microplate reader (TECAN). Nitrite concentrations were calculated from a sodium nitrite (NaNO₂) standard curve and expressed as μM nitrite. All samples were analyzed in triplicates, and results are presented as the mean ± SEM of at least three independent biological experiments.

### Flow Cytometry Analysis of Macrophage Surface Markers

RAW264.7 macrophages were harvested using Cell Dissociation Buffer (Sigma-Aldrich, Cat. # T4049) for 5 min at 37°C and gently scraped to obtain a single-cell suspension. Cells were washed twice with PBS and incubated with anti-mouse Fc receptor blocking antibody for 10 min on ice. Cells were then stained with fluorochrome-conjugated antibodies against CD80, CD86, and MHC class II (I-A/I-E) for 30-45 min on ice in the dark. Antibody details are provided in Table 2. Following staining, cells were washed twice and analyzed using a BD FACSCanto II flow cytometer. Single-color controls were used for compensation, and unstained cells were used to establish background fluorescence. Data were analyzed using FACSDiva and FlowJo software. Cells were gated based on forward and side-scatter characteristics, followed by doublet exclusion. Results are presented as the percentage of positive cells or mean fluorescence intensity (MFI), as indicated in the figure legends.

### Flow Cytometric Analysis of Mitochondrial and ROS Parameters

For analysis of mitochondrial parameters and intracellular ROS, macrophages were incubated with MitoTracker Green FM (50 nM; Thermo Fisher Scientific, Cat. #M7514), TMRE (150 nM; Thermo Fisher Scientific, Cat. #T669), MitoSOX Red (5 μM; Thermo Fisher Scientific, Cat. #M36008), or DCFDA (10 μM; Sigma-Aldrich, Cat. #35845) for 30 min at 37°C in the dark, according to the manufacturers’ recommendations. MitoTracker Green was used to assess mitochondrial content, TMRE to measure mitochondrial membrane potential, MitoSOX Red to measure mitochondrial superoxide, and DCFDA to measure intracellular ROS. Cells were washed twice with pre-warmed PBS and immediately analyzed using a BD FACSCanto II flow cytometer. A minimum of 10,000-20,000 events was acquired per sample. Data were analyzed using FACSDiva and FlowJo software. Results are presented as MFI, or as TMRE fluorescence normalized to MitoTracker Green fluorescence where indicated.

### Immunofluorescence Microscopy

RAW264.7 cells or bone marrow-derived macrophages (BMDMs) on glass coverslips were fixed with 4% paraformaldehyde (15 min), permeabilized with 0.1% Triton X-100 (10 min), and blocked with 2% BSA (1 h). Samples were incubated overnight at 4°C with primary antibodies (Table 2), followed by fluorophore-conjugated secondary antibodies (1 h) protected from light. For actin and nuclear visualization, cells were stained with Phalloidin-Tetramethyl-Rhodamine B isothiocyanate (Cat. #P1951, Sigma-Aldrich) and DAPI (Cat. #10236276001, Roche), respectively. Where indicated, live cells were pre-stained with Mito Tracker™ Red (100 nM, 30 min; Cat. #M22426, Thermo Fisher Scientific) prior to fixation. Coverslips were mounted using ProLong™ Gold Antifade (Cat. #P36934, Invitrogen). Confocal images were acquired on an Olympus FV3000 or Nikon-SIM microscope using a 63X oil-immersion objective with identical settings across experimental groups. Quantification was performed using Fiji/ImageJ.

### Transmission Electron Microscopy (TEM)

Macrophage cells were collected and suspended in phosphate buffer (PB), followed by centrifugation to obtain a cell pellet. The pellets were then immersed in TEM fixative and maintained at 4 °C for 2 h. After fixation, the samples were centrifuged, the fixative was discarded, and the pellets were washed and resuspended in PB. The specimens were subsequently processed for transmission electron microscopy (TEM), and ultrathin sections approximately 60-90 nm thick were obtained. The sections were contrasted sequentially with uranyl acetate and alkaline lead citrate before imaging. TEM observations were performed on a JEM-1400 Flash transmission electron microscope equipped with a tungsten filament electron source and a high-sensitivity sCMOS camera. Images were collected at an accelerating voltage of 120 kV and a magnification of 12,000X.

### Cellular and Mitochondrial Morphology Quantification

Cellular morphology was quantified from confocal micrographs based on phalloidin staining. Images were background-subtracted and thresholded to define individual cell boundaries. Cell area, length, perimeter, circularity, aspect ratio, and solidity were measured using the Particle Analysis function in ImageJ. Mitochondrial morphology was quantified from high-resolution confocal Z-stack images of Mito Tracker-stained cells using ImageJ with the Mitochondria Analyzer plugin. Images were subjected to background correction, noise reduction, and thresholding to segment mitochondrial structures. Mitochondrial network features, including the number of branches and junctions, total and mean branch length, and network connectivity, were quantified from individual cells. Measurements were obtained from the indicated number of cells per experimental condition, as specified in the figure legends.

### Mitochondrial Isolation

Mitochondria were isolated from untreated (M0) and IFN-γ stimulated (M1) RAW264.7 macrophages (4 × 10⁷ cells per sample) using the Mitochondria Isolation Kit for Cultured Cells (Cat. #HY-K1060, Med Chem Express, USA) according to the manufacturer’s instructions. Briefly, cells were harvested, washed with ice-cold PBS, and homogenized in the supplied mitochondrial isolation buffer using a Dounce homogenizer on ice. The homogenate was subjected to differential centrifugation to separate the mitochondrial and cytosolic fractions according to the manufacturer’s protocol. Protein concentrations of the isolated mitochondrial fractions were determined using the Bio-Red protein Assay Dye Reagent. Equal amounts of protein were analyzed by Western blotting to determine IIGP1 localization. Voltage-dependent anion channel (VDAC) was used as a mitochondrial marker, whereas β-actin served as a cytosolic marker to assess the purity of the isolated fractions.

### Measurement of Oxygen Consumption Rate (OCR) and Extracellular Acidification Rate (ECAR)

Cellular metabolic function was assessed by measuring the oxygen consumption rate (OCR) and extracellular acidification rate (ECAR) using a Seahorse XF24 Extracellular Flux Analyzer (Agilent Technologies). RAW264.7 macrophages were seeded at 2 × 10⁴ cells per well in Seahorse XF24 cell culture microplates and allowed to adhere overnight. Cells were polarized under the indicated experimental conditions for 24 h prior to metabolic analysis. On the day of the assay, culture medium was replaced with pre-warmed Seahorse XF assay medium, and cells were incubated for 1 h at 37°C in a non-CO₂ incubator to allow temperature and pH equilibration. For the mitochondrial stress test, OCR was measured under basal conditions followed by sequential injections of oligomycin (1 μM), FCCP (1 μM), and rotenone (0.5 μM). Basal respiration, ATP-linked respiration, maximal respiratory capacity, spare respiratory capacity, and coupling efficiency were calculated using Wave software (Agilent Technologies) according to the manufacturer’s instructions. For the glycolysis stress test, ECAR was measured following sequential injections of glucose (10 mM), oligomycin (1 μM), and 50 mM 2-deoxy-D-glucose (2-DG) to determine glycolysis, glycolytic capacity, and glycolytic reserve. Data were analyzed using Wave software (Agilent Technologies) and are presented as the mean of at least three independent biological experiments.

### In-solution digestion and LFQ analysis

The cell pellet of differentiated (M0) and polarized RAW 264.7 macrophages (M1-IFN-γ and M2-IL-4) were collected by gentle scraping of the cells with a cell scraper, followed by PBS washing and finally a spun at 2000 rpm for 10 mins. The cell pellet was then lysed using urea lysis buffer and passed through a 3 kDa filter column. The protein content was measured by Bradford assay prior to use. Twenty-five micrograms of protein from each sample was reduced (10 mM DTT), alkylated (50mM IAA) and quenched (10mM DTT). This was followed by protein digestion using overnight trypsin treatment (1μg) in a 37°C water bath. The digested samples were dried in SpeedVac and then reconstituted in 25μl 0.1%TFA. 0.5μl β-galactosidase was added to each sample as an internal standard. The tryptic digest was then desalted using Zip-Tip (Merck Millipore) and eluted in 50% ACN + 0.1%TFA (50:50). The fragmented protein was then dried and resuspended in 0.1% formic acid. The peptide concentration was estimated using nano drop (Thermo-Fisher Scientific), and the samples were injected into LC/MS (Orbitrap Fusion mass spectrometer, Thermo-Fisher Scientific). The Raw data from MS were analyzed for differential expression of protein using Proteome Discoverer 2.2 (Thermo-Fisher Scientific).

### Immunoprecipitation coupled with mass spectrometry (IP-MS)

To identify proteins interacting with IIGP1, immunoprecipitation coupled with mass spectrometry (IP-MS) was performed using lysates prepared from RAW 264.7 cells. Cells were cultured and stimulated with IFN-γ for 24 hrs before harvesting. Cell pellets were lysed in NETN buffer (20 mM Tris-HCl, pH 8.0, 100 mM NaCl, 1 mM EDTA, and 0.5% NP-40) supplemented with protease and phosphatase inhibitor cocktails. The lysates were clarified by centrifugation at 16,000 × g for 20 min at 4°C and precleared with 30 μL of Protein A/G magnetic beads (Cat. #HY-K1060, Med Chem Express, USA). For immunoprecipitation, 5 μg of rabbit polyclonal anti-IIGP1 antibody (Table 2) or rabbit IgG Isotype control (Cat. #02-6102, Invitrogen, USA) was coupled to 50 μL of Protein A/G magnetic beads and incubated with 1.5 mg of precleared protein lysate overnight at 4°C with gentle rotation. Following incubation, the beads were washed sequentially with PBST and ammonium bicarbonate buffer to remove nonspecifically bound proteins. Bound proteins were eluted in 0.1% RapiGest containing 5 mM DTT and digested overnight with 2 μg of sequencing-grade trypsin. The resulting peptides were desalted using C18 Zip-Tips and analyzed by liquid chromatography-tandem mass spectrometry (LC-MS/MS) on an Orbitrap Fusion mass spectrometer (Thermo Fisher Scientific). Mass spectrometry data were processed using Proteome Discoverer version 2.2, with protein identification filtered at a 1% false discovery rate (FDR) and requiring a minimum of two unique peptides per protein.

### Bacterial Phagocytosis and Intracellular Killing Assays

Macrophages were stimulated with IFN-γ (100 ng/mL) for 24 h before infection. For intracellular bacterial survival assays, cells were infected with *Escherichia coli* DH5α or *Salmonella Typhimurium* at an MOI of 1:40 for 1 h. Cells were washed with PBS and incubated with complete medium containing gentamicin (100 μg/mL; Cat. #15750078, Gibco) for an additional 1 h. The medium was then replaced with maintenance medium containing gentamicin (10 μg/mL), and cells were incubated for the indicated times. At each time point, cells were washed with PBS and lysed with 0.1% Triton X-100. Serial dilutions of the lysates were plated on LB agar and incubated overnight at 37 °C. Intracellular bacterial burden was determined by enumeration of colony-forming units (CFU). For bacterial uptake analysis, IFN-γ-stimulated macrophages were infected with mCherry-expressing *S. Typhimurium* at an MOI of 1:40 for 1 h. Cells were washed three times with PBS to remove extracellular bacteria and fixed with 4% paraformaldehyde. Nuclei were stained with DAPI, and images were acquired using an Olympus FV3000 laser-scanning confocal microscope. Bacterial uptake was quantified by measuring intracellular mCherry fluorescence using ImageJ.

### STRING Database Analysis

Protein-protein interaction (PPI) networks and functional associations were analyzed using the STRING database (version 12.5) (http://string.embl.de/). To identify interacting clusters, k-means clustering was implemented within the STRING platform based on evidence scores, applying a minimum required interaction score (confidence cutoff) of 0.40. (Optional addition if enrichment was also run: Functional enrichment analyses, including Gene Ontology (GO) and pathway analyses, were subsequently performed on the clustered networks).

### Promoter Analysis

Putative transcription factor binding sites within the Mus musculus Iigp1 promoter (NCBI Reference Sequence: NM_021792) were identified using the Eukaryotic Promoter Database (EPD) (Swiss Institute of Bioinformatics). The promoter region spanning −2000 bp to +100 bp relative to the transcription start site (TSS) was analyzed using the JASPAR CORE 2018 vertebrate transcription factor motif database. Transcription factor binding motifs were predicted using a significance threshold of P < 0.001, and putative STAT1 binding sites within the Iigp1 promoter were identified for further analysis.

### Chromatin Immunoprecipitation Sequencing (ChIP-seq) Analysis

To analyze STAT1 binding dynamics at the Iigp1 promoter locus, public functional genomics data were leveraged from the NCBI Gene Expression Omnibus (GEO) repository under the dataset series accession number *GSE33913*. The continuous enrichment data track capturing genomic STAT1 recruitment corresponds to sample *GSM838683*, representing wild-type (WT) bone marrow-derived macrophages (BMDMs) stimulated with IFN-γ for 6 hours. Raw signal density tracks and peak boundaries were visualized and mapped using the UCSC Genome Browser on the mouse reference genome assembly NCBI37/mm9.

### Statistical analyses

Quantitative data are presented as the mean ± standard error of the mean (SEM), unless indicated otherwise, from independent experiments as indicated in the figure legends. Statistical analyses were performed using GraphPad Prism 9. The specific tests used are indicated in the figure legends. Statistical significance is denoted by asterisks: ns p > 0.05, *p < 0.05, **p < 0.01, ***p <0.001, and ****p < 0.0001.

### AI Tools

Artificial intelligence (AI) tools were used solely for language editing, grammatical correction, and textual refinement of the manuscript. AI tools were not used for experimental design, data acquisition, data analysis, statistical analysis, scientific interpretation, or figure generation. All experimental work, data analysis and interpretation, figures, and scientific conclusions were performed and developed by the authors.

## Supporting information

https://drive.google.com/drive/u/0/folders/1n3tW44xjkQWHWoLwYU1_Jg9zfknWdtKp

## Author statement

**Arvind Sahu and Anup Kumar Singh**: Conceptualization, Methodology. **Anup Kumar Singh**: Formal analysis, Writing Original draft preparation. **Anup Kumar Singh**: Investigation. **Arvind Sahu**: Supervision, Funding acquisition. **Arvind Sahu, Manas Kumar Santra, Srikanth Repole, Rajender K Motiani**, **Samriddhi Sharma**, **Kriti Ahuja**: Writing, Reviewing and Editing.

## Funding and additional information

This work was supported by a research grant from the Anusandhan National Research Foundation (CRG/2023/001054), Ministry of Science and Technology, Government of India (to A.S.), Core funding from NCCS, and J. C. Bose National Fellowship (to A.S.)

## Declaration of interests

The authors declare that they do not have any conflicts of interest related to this study.

## Acknowledgement

We express our gratitude to Dr. Jomon Joseph (NCCS, Pune) for expert advice on CRISPR-Cas9 genome editing, and Dr. Chittur V. Srikanth (RCB, Faridabad) for kindly providing the Salmonella strains. We acknowledge the support and use of the proteomics, microscopy, and bioimaging facilities at NCCS. This study forms part of the Ph.D. thesis of A.K.S., registered at Savitribai Phule Pune University. A.K.S., S.S., and K.A. acknowledge the receipt of doctoral research fellowships from CSIR, ICMR, and DBT, New Delhi, respectively.

## Abbreviations

IFN-γ: Interferon-γ
TNF-α: tumor necrosis factor-α
LPS: lipopolysaccharide
iNOS: inducible nitric oxide synthase
ROS: Reactive Oxygen Species
RNS: Reactive Nitrogen Species
BMDM: Bone-Marrow Derived Macrophage
Mitoch: ondrial Reactive Oxygen Species

## Supporting information

**Supplementary Figure 1. Quantitative morphometric analysis of macrophage activation states. (A)** Workflow of morphology analysis of macrophages using QuPath and ImageJ software. **(B-E)** Confocal microscopy and high-throughput morphological quantification of **(B, C)** bone marrow-derived macrophages (BMDMs) and **(D, E)** RAW264.7 cells. Macrophages were polarized toward M1 (IFN-γ) or M2 (IL-4) states for 24 h, fixed with 4% paraformaldehyde, and counterstained with DAPI to visualize nuclei (blue) alongside state-specific phalloidin conjugates to visualize the F-actin cytoskeleton (Alexa Fluor™ 488-phalloidin [green] for BMDMs; Rhodamine-phalloidin [red] for RAW264.7 cells. Cell boundaries from at least 300 individual cells per macrophage phenotype were manually annotated using QuPath (v0.3.2) and ImageJ. Quantified geometric parameters include total cell area, cellular perimeter, major axis length, and minor axis length. Structural geometry and cell elongation profiles were mathematically derived using aspect ratio, circularity, and eccentricity metrics. Data represent mean ± SD (n=3 independent biological replicates). Statistical significance was determined by one-way ANOVA with Tukey’s post-hoc test (*p* ≤ is less than or equal to ≤ 0.05, ** *p* ≤ is less than or equal to ≤ 0.01, *** *p* ≤ is less than or equal to ≤ 0.001, **** *p* ≤ is less than or equal to ≤ 0.0001).

**Supplementary Figure 2. Transcriptional and proteomic validation of M1 and M2 macrophage polarization states.** Transcriptional profiling of state-specific polarization markers in primary bone marrow-derived macrophages (BMDMs). Cells were left unstimulated (M0), stimulated with IFN-γ (100 ng/mL, M1), or stimulated with IL-4 (20 ng/mL, M2) for 24 h. Quantitative RT-PCR analysis was performed to evaluate the expression of **(A)** classical M1-associated transcripts (*Nos2*, *Il6*, *Il1b*) and **(B)** M2-associated transcripts (*Arg1*, *Ym-1* and *FIZZ-1*]). Transcript levels are presented relative to M0 baseline controls. **(C, D)** Transcriptional profiling of polarization markers in RAW264.7 cells under identical M0, M1, and M2 polarizing conditions. Quantitative RT-PCR was used to assess **(C)** M1-associated transcripts (*Nos2*, *Il6*, *Il1b*) and **(D)** M2-associated transcripts (*Arg1* and *Mrc1*). **(E, F)** Western blot validation of polarization markers in **(E)** primary BMDMs and **(F)** RAW264.7 cells. Whole-cell lysates harvested from M0, M1, and M2 macrophages were analyzed by immunoblotting using anti-iNOS (M1 marker) and anti-Arg-1 (M2 marker) antibodies, with β-actin serving as the loading control. Data represent mean ± SD (n=3 independent biological replicates). Statistical significance was determined by one-way ANOVA with Tukey’s post-hoc test (p ≤ is less than or equal to ≤ 0.05, ** p ≤ is less than or equal to ≤ 0.01, *** p ≤ is less than or equal to ≤ 0.001, **** p ≤ is less than or equal to ≤ 0.0001).

**Supplementary figure 3: Culture medium acidification in WT and IIGP1-KO macrophages following polarization. (A)** RAW264.7 macrophages were seeded in 6-well plates (1.5 × 10⁶ cells per well) polarized toward M0, M1, or M2 phenotypes for 24 h using state-specific cytokine stimuli. Representative culture vessel photographs depict the macroscopic color changes of the phenol red-containing culture medium, alongside a corresponding quantification plot displaying the exact pH values of the harvested culture supernatants across each genotype and polarization state. Data are presented as mean ± SD from three independent biological replicates. Statistical significance was determined by one-way ANOVA with Tukey’s post-hoc test (p ≤ is less than or equal to ≤ 0.05, ** p ≤ is less than or equal to ≤ 0.01, *** p ≤ is less than or equal to ≤ 0.001, **** p ≤ is less than or equal to ≤ 0.0001).

## Notes

### Competing Interest Statement

The authors have declared no competing interest.

## References

1. Guan, F., Wang, R., Yi, Z., Luo, P., Liu, W., Xie, Y., Liu, Z., Xia, Z., Zhang, H., & Cheng, Q. (2025). Tissue macrophages: origin, heterogenity, biological functions, diseases and therapeutic targets. Signal transduction and targeted therapy, 10(1), 93. 10.1038/s41392-025-02124-y

2. Hirayama, D., Iida, T., & Nakase, H. (2017). The Phagocytic Function of Macrophage-Enforcing Innate Immunity and Tissue Homeostasis. International journal of molecular sciences, 19(1), 92. 10.3390/ijms19010092

3. Wynn, T. A., & Vannella, K. M. (2016). Macrophages in Tissue Repair, Regeneration, and Fibrosis. Immunity, 44(3), 450–462. 10.1016/j.immuni.2016.02.015

4. Lendeckel, U., Venz, S., & Wolke, C. (2022). Macrophages: shapes and functions. Chemtexts, 8(2), 12. 10.1007/s40828-022-00163-4

5. Murray, P. J., & Wynn, T. A. (2011). Protective and pathogenic functions of macrophage subsets. Nature reviews. Immunology, 11(11), 723–737. 10.1038/nri3073

6. Murray P. J. (2017). Macrophage Polarization. Annual review of physiology, 79, 541–566. 10.1146/annurev-physiol-022516-034339

7. Stein M. Interleukin 4 potently enhances murine macrophage mannose receptor activity: a marker of alternative immunologic macrophage activation. J Exp Med. (1992) 176:287–92. doi: 10.1084/jem.176.1.287

8. Doyle AG, Herbein G, Montaner LJ, Minty AJ, Caput D, Ferrara P, et al. Interleukin-13 alters the activation state of murine macrophages in vitro: Comparison with interleukin-4 and interferon-γ. Eur J Immunol. (1994) 24:1441–5. doi: 10.1002/eji.183024063

9. Mantovani A, Biswas SK, Galdiero MR, Sica A, Locati M. Macrophage plasticity and polarization in tissue repair and remodelling. J Pathol. (2013) 229:176–85. doi: 10.1002/path.4133

10. Viola, A., Munari, F., Sánchez-Rodríguez, R., Scolaro, T., & Castegna, A. (2019). The Metabolic Signature of Macrophage Responses. Frontiers in immunology, 10, 1462. 10.3389/fimmu.2019.01462

11. O’Neill, L. A., & Pearce, E. J. (2016). Immunometabolism governs dendritic cell and macrophage function. The Journal of experimental medicine, 213(1), 15–23. 10.1084/jem.20151570

12. Freemerman, A. J., Johnson, A. R., Sacks, G. N., Milner, J. J., Kirk, E. L., Troester, M. A., Macintyre, A. N., Goraksha-Hicks, P., Rathmell, J. C., & Makowski, L. (2014). Metabolic reprogramming of macrophages: glucose transporter 1 (GLUT1)-mediated glucose metabolism drives a proinflammatory phenotype. The Journal of biological chemistry, 289(11), 7884–7896. 10.1074/jbc.M113.522037

13. Batista-Gonzalez, A., Vidal, R., Criollo, A., & Carreño, L. J. (2020). New Insights on the Role of Lipid Metabolism in the Metabolic Reprogramming of Macrophages. Frontiers in immunology, 10, 2993. 10.3389/fimmu.2019.02993

14. Tur, J., Pereira-Lopes, S., Vico, T., Marín, E. A., Muñoz, J. P., Hernández-Alvarez, M., Cardona, P. J., Zorzano, A., Lloberas, J., & Celada, A. (2020). Mitofusin 2 in Macrophages Links Mitochondrial ROS Production, Cytokine Release, Phagocytosis, Autophagy, and Bactericidal Activity. Cell reports, 32(8), 108079. 10.1016/j.celrep.2020.108079

15. Wu, D., Wang, Y., Hu, J., Xu, Y., Gong, D., Wu, P., Dong, J., He, B., Qian, H., & Wang, G. (2023). Rab26 promotes macrophage phagocytosis through regulation of MFN2 trafficking to mitochondria. The FEBS journal, 290(16), 4023–4039. 10.1111/febs.16793

16. Lloberas, J., Muñoz, J. P., Hernández-Álvarez, M. I., Cardona, P. J., Zorzano, A., & Celada, A. (2020). Macrophage mitochondrial MFN2 (mitofusin 2) links immune stress and immune response through reactive oxygen species (ROS) production. Autophagy, 16(12), 2307–2309. 10.1080/15548627.2020.1839191

17. Van den Bossche, J., Baardman, J., Otto, N. A., van der Velden, S., Neele, A. E., van den Berg, S. M., Luque-Martin, R., Chen, H. J., Boshuizen, M. C., Ahmed, M., Hoeksema, M. A., de Vos, A. F., & de Winther, M. P. (2016). Mitochondrial Dysfunction Prevents Repolarization of Inflammatory Macrophages. Cell reports, 17(3), 684–696. 10.1016/j.celrep.2016.09.008

18. Tannahill, G. M., Curtis, A. M., Adamik, J., Palsson-McDermott, E. M., McGettrick, A. F., Goel, G., Frezza, C., Bernard, N. J., Kelly, B., Foley, N. H., Zheng, L., Gardet, A., Tong, Z., Jany, S. S., Corr, S. C., Haneklaus, M., Caffrey, B. E., Pierce, K., Walmsley, S., Beasley, F. C., … O’Neill, L. A. (2013). Succinate is an inflammatory signal that induces IL-1β through HIF-1α. Nature, 496(7444), 238–242. 10.1038/nature11986

19. Wang, Y., Li, N., Zhang, X., & Horng, T. (2021). Mitochondrial metabolism regulates macrophage biology. The Journal of biological chemistry, 297(1), 100904. 10.1016/j.jbc.2021.100904

20. Li, P., Wang, G., Peng, Z., Zhang, L., Yang, F., Wei, Y., Pan, M., Zang, H., & Zhou, M. (2025). The Role of Mitochondrial Regulation in Macrophage Polarization: Implications for the Pathogenesis of Rheumatoid Arthritis. Journal of inflammation research, 18, 17163–17183. 10.2147/JIR.S560635

21. Kim, M. E., Lim, Y., & Lee, J. S. (2025). Mitochondrial Dysfunction and Metabolic Reprogramming in Chronic Inflammatory Diseases: Molecular Insights and Therapeutic Opportunities. Current issues in molecular biology, 47(12), 1042. 10.3390/cimb47121042

22. Ji, F., Zhang, L., Ning, L., Zhang, M., & Zhang, J. (2026). Crosstalk of mitochondrial dysfunction and macrophage polarization in sepsis. Frontiers in immunology, 17, 1692597. 10.3389/fimmu.2026.1692597

23. Liu, J., Wang, M., & Zhao, Y. (2025). The Regulatory Network of Transcription Factors in Macrophage Polarization. ImmunoTargets and therapy, 14, 555–575. 10.2147/ITT.S494550

24. Jha, A. K., Huang, S. C., Sergushichev, A., Lampropoulou, V., Ivanova, Y., Loginicheva, E., Chmielewski, K., Stewart, K. M., Ashall, J., Everts, B., Pearce, E. J., Driggers, E. M., & Artyomov, M. N. (2015). Network integration of parallel metabolic and transcriptional data reveals metabolic modules that regulate macrophage polarization. Immunity, 42(3), 419–430. 10.1016/j.immuni.2015.02.005

25. Michalska, A., Blaszczyk, K., Wesoly, J., & Bluyssen, H. A. R. (2018). A Positive Feedback Amplifier Circuit That Regulates Interferon (IFN)-Stimulated Gene Expression and Controls Type I and Type II IFN Responses. Frontiers in immunology, 9, 1135. 10.3389/fimmu.2018.01135

26. Pawlowski, N. (2010). Dynamin self-assembly and the vesicle scission mechanism: how dynamin oligomers cleave the membrane neck of clathrin-coated pits during endocytosis. BioEssays: news and reviews in molecular, cellular and developmental biology, 32 *12*, 1033–9.

27. Taylor, G. A., Collazo, C. M., Yap, G. S., Nguyen, K., Gregorio, T. A., Taylor, L. S., Eagleson, B., Secrest, L., Southon, E. A., Reid, S. W., Tessarollo, L., Bray, M., McVicar, D. W., Komschlies, K. L., Young, H. A., Biron, C. A., Sher, A., & Vande Woude, G. F. (2000). Pathogen-specific loss of host resistance in mice lacking the IFN-gamma-inducible gene IGTP. Proceedings of the National Academy of Sciences of the United States of America, 97(2), 751–755. 10.1073/pnas.97.2.751

28. Yamamoto, M., Okuyama, M., Ma, J. S., Kimura, T., Kamiyama, N., Saiga, H., Ohshima, J., Sasai, M., Kayama, H., Okamoto, T., Huang, D. C., Soldati-Favre, D., Horie, K., Takeda, J., & Takeda, K. (2012). A cluster of interferon-γ-inducible p65 GTPases plays a critical role in host defense against Toxoplasma gondii. Immunity, 37(2), 302–313. 10.1016/j.immuni.2012.06.009

29. Collazo, C. M., Yap, G. S., Sempowski, G. D., Lusby, K. C., Tessarollo, L., Vande Woude, G. F., Sher, A., & Taylor, G. A. (2001). Inactivation of LRG-47 and IRG-47 reveals a family of interferon gamma-inducible genes with essential, pathogen-specific roles in resistance to infection. The Journal of experimental medicine, 194(2), 181–188. 10.1084/jem.194.2.1814

30. Kim, B. H., Shenoy, A. R., Kumar, P., Bradfield, C. J., & MacMicking, J. D. (2012). IFN-inducible GTPases in host cell defense. Cell host & microbe, 12(4), 432–444. 10.1016/j.chom.2012.09.007

31. Coers J. (2013). Self and non-self-discrimination of intracellular membranes by the innate immune system. PLoS pathogens, 9(9), e1003538. 10.1371/journal.ppat.1003538

32. Al-Zeer, M. A., Al-Younes, H. M., Braun, P. R., Zerrahn, J., & Meyer, T. F. (2009). IFN-gamma-inducible Irga6 mediates host resistance against Chlamydia trachomatis via autophagy. PloS one, 4(2), e4588. 10.1371/journal.pone.0004588

33. Ferreira-da-Silva, M.daF., Springer-Frauenhoff, H. M., Bohne, W., & Howard, J. C. (2014). Identification of the microsporidian Encephalitozoon cuniculi as a new target of the IFNγ-inducible IRG resistance system. PLoS pathogens, 10(10), e1004449. 10.1371/journal.ppat.1004449

34. Kuriakose, T., & Kanneganti, T. D. (2017). War on viruses: LC3 recruits GTPases. Cell Host & Microbe, 22(1), 7–9.

35. Biering, S. B., Choi, J., Halstrom, R. A., Brown, H. M., Beatty, W. L., Lee, S., McCune, B. T., Dominici, E., Williams, L. E., Orchard, R. C., Wilen, C. B., Yamamoto, M., Coers, J., Taylor, G. A., & Hwang, S. (2017). Viral Replication Complexes Are Targeted by LC3-Guided Interferon-Inducible GTPases. Cell host & microbe, 22(1), 74–85.e7. 10.1016/j.chom.2017.06.005

36. Singh, S. B., Ornatowski, W., Vergne, I., Naylor, J., Delgado, M., Roberts, E., Ponpuak, M., Master, S., Pilli, M., White, E., Komatsu, M., & Deretic, V. (2010). Human IRGM regulates autophagy and cell-autonomous immunity functions through mitochondria. Nature cell biology, 12(12), 1154–1165. 10.1038/ncb2119

37. Mills C. D. (2015). Anatomy of a discovery: m1 and m2 macrophages. Frontiers in immunology, 6, 212. 10.3389/fimmu.2015.00212

38. Martens S, Sabel K, Lange R, Uthaiah R, Wolf E, et al. (2004) Mechanisms regulating the positioning of mouse p47 resistance GTPases LRG-47 and IIGP1 on cellular membranes: retargeting to plasma membrane induced by phagocytosis. J Immunol 173: 2594–2606.

39. Henry, S. C., Traver, M., Daniell, X., Indaram, M., Oliver, T., & Taylor, G. A. (2010). Regulation of macrophage motility by Irgm1. Journal of leukocyte biology, 87(2), 333–343. 10.1189/jlb.0509299

40. Traver, M. K., Henry, S. C., Cantillana, V., Oliver, T., Hunn, J. P., Howard, J. C., Beer, S., Pfeffer, K., Coers, J., & Taylor, G. A. (2011). Immunity-related GTPase M (IRGM) proteins influence the localization of guanylate-binding protein 2 (GBP2) by modulating macroautophagy. The Journal of biological chemistry, 286(35), 30471–30480. 10.1074/jbc.M111.251967

41. Zerrahn, J., Schaible, U. E., Brinkmann, V., Guhlich, U., & Kaufmann, S. H. (2002). The IFN-inducible Golgi- and endoplasmic reticulum-associated 47-kDa GTPase IIGP is transiently expressed during listeriosis. *Journal of immunology (Baltimore*, Md.: 1950), *168*(7), 3428–3436. 10.4049/jimmunol.168.7.3428

42. Park, S., Choi, J., Biering, S. B., Dominici, E., Williams, L. E., & Hwang, S. (2016). Targeting by AutophaGy proteins (TAG): Targeting of IFNG-inducible GTPases to membranes by the LC3 conjugation system of autophagy. Autophagy, 12(7), 1153–1167. 10.1080/15548627.2016.1178447

43. Henry, S. C., Schmidt, E. A., Fessler, M. B., & Taylor, G. A. (2014). Palmitoylation of the immunity related GTPase, Irgm1: impact on membrane localization and ability to promote mitochondrial fission. PloS one, 9(4), e95021.

44. McWhorter, F. Y., Wang, T., Nguyen, P., Chung, T., & Liu, W. F. (2013). Modulation of macrophage phenotype by cell shape. Proceedings of the National Academy of Sciences of the United States of America, 110(43), 17253–17258. 10.1073/pnas.1308887110

45. Sharma, P., Vijaykumar, A., Raghavan, J. V., Rananaware, S. R., Alakesh, A., Bodele, J., Rehman, J. U., Shukla, S., Wagde, V., Nadig, S., Chakrabarti, S., Visweswariah, S. S., Nandi, D., Gopal, B., & Jhunjhunwala, S. (2022). Particle uptake driven phagocytosis in macrophages and neutrophils enhances bacterial clearance. Journal of controlled release: official journal of the Controlled Release Society, 343, 131–141. 10.1016/j.jconrel.2022.01.030

46. Depierre, M., Mularski, A., Ruppel, A., Le Clainche, C., Balland, M., & Niedergang, F. (2025). A crosstalk between adhesion and phagocytosis integrates macrophage functions into their microenvironment. iScience, 28(4), 112067. 10.1016/j.isci.2025.112067

47. Diotallevi, M., Outeiral, C., Patel, P., Purvis, G. S. D., Chuaiphichai, S., Nicol, T., Ayaz, F., Nissley, D. A., Krasnoselska, G. O., Hester, S., McVey, J. H., Fischer, R., Kessler, B., Deane, C. M., Owens, R. J., Channon, K. M., & Crabtree, M. J. (2026). iNOS modulates inflammatory responses in an NO-independent manner through direct interaction with IRG1 in mitochondria. Nature metabolism, 8(4), 855–868. 10.1038/s42255-026-01492-1

48. Bailey, J. D., Diotallevi, M., Nicol, T., McNeill, E., Shaw, A., Chuaiphichai, S., Hale, A., Starr, A., Nandi, M., Stylianou, E., McShane, H., Davis, S., Fischer, R., Kessler, B. M., McCullagh, J., Channon, K. M., & Crabtree, M. J. (2019). Nitric Oxide Modulates Metabolic Remodeling in Inflammatory Macrophages through TCA Cycle Regulation and Itaconate Accumulation. Cell reports, 28(1), 218–230.e7. 10.1016/j.celrep.2019.06.018

