## Supplementary material for "The IFN-γ-inducible GTPase IIGP1 regulates mitochondrial remodeling and immunometabolic reprogramming during inflammatory macrophage polarization": https://drive.google.com/drive/u/0/folders/1n3tW44xjkQWHWoLwYU1_Jg9zfknWdtKp: Main Table.docx.pdf

Table-1: **Primer sequences used for qRT-PCR**

| **Primers** | **Forward (5'-3')** | **Reverse (5'-3')** |
|---|---|---|
| mβactin | AGCCATGTACGTAGCCATCC | CTCTCAGCTGTGGTGGTGAA |
| iNOS | ACATCGACCCGTCCACAGTAT | CAGAGGGGTAGGCTTGTCTC |
| IL-6 | ACAACCACGGCCTTCCCTACTT | CACGATTTCCCAGAGAACATGTG |
| IL-1β | GAAATGCCACCTTTTGACAGTC | CTGGATGCTCTCATCAGGACA |
| Arg1 | CTCCAAGCCAAAGTCCTTAGAG | GGAGCTGTCATTAGGGACATCA |
| Ccl2 | CAGCCAGATGCAGTTAACGC | GCCTACTCATTGGGATCATCTTG |
| Ym1 | CATGAGCAAGACTTGCGTGAC | GGTCCAAACTTCCATCCTCCA |
| Mrc1 | CTCTGTTCAGCTATTGGACGC | TGGCACTCCCAAACATAATTTGA |
| FIZZ1 | TCCAGCTAACTATCCCTCCACTGT | GGCCCATCTGTTCATAGTCTTGA |
| IIGP1 | AATACCTGCCTCACGCTCAT | GCTACTCTGTGGGTTCTGGC |

Table 2: **Antibodies used for western blots**

| **Antibody** | **Concentration** | **Species** | **Product No.** |
|---|---|---|---|
| IIGP1 | 1:1000 | Rabbit | Abclonal #A15226 |
| iNOS (D6B6S) | 1:1000 | Rabbit | Cell Signal #13150S |
| STAT1 | 1:1000 | Rabbit | Cell Signal #9172S |
| P-Stat1 (Y701) | 1:1000 | Rabbit | Cell Signal #9167S |
| OPA-1 (D6U6N) | 1:1500 | Rabbit | Cell Signal # 80471 |
| Drp-1 (D6C7) | 1:1000 | Rabbit | Cell Signal # 8570 |
| β-Actin | 1:1000 | Mouse | Abclonal #AC004 |
| VDAC | 1:1000 | Rabbit | Cell Signal # 4866 |
| Anti-rabbit IgG (H+L)-HRP Conjugate | 1:10000 | Goat | Bio-Rad # 1706515 |
| Anti-mouse IgG (H+L)-HRP Conjugate | 1:10000 | Goat | Bio-Rad #172-1034 |
| Anti-rabbit IgG Alexa Fluor 488-conjugated | 1:1000 | Goat | Invitrogen # A-11008 |
| CD86 | 1:1000 | Rabbit | Bio Legend # 105021 |
| MHC-II (I-A/I-E) | 1:1000 | Rabbit | Bio Legend #107612 |
| CD80 | 1:1000 | Rabbit | Bio Legend #123109 |
| CD16/32 (Fc-Block) | 1:100 | Rabbit | BD Bioscience # 553141 |

Table-3: **Primer sequences used for CRISPR-Cas9-sg-Iigp1**

| **sgRNA** | **Forward (5'-3')** | **Reverse (5'-3')** |
|---|---|---|
| **Iigp1_sgRNA1** | GATCCG**ACACCCCAATATACCC**[illegible]**G**GTTTTAGAGCTAGAAATAGCAA[illegible]AAAATAAGGCTAGTCCGTTATCA[illegible]TGAAAAAGTGGCACCGAGTCGG[illegible]TTTTTT | CTAGAAAAAAGCACCGACTCGGTGCCACTTTTTCAAGTTGATAACGGACTAGCCTTATTTTAACTTGCTATTTCTAGCTCTAAAACCATTGGGTATATTGGGGTGTCG |
