## Supplementary figures and images for "The IFN-γ-inducible GTPase IIGP1 regulates mitochondrial remodeling and immunometabolic reprogramming during inflammatory macrophage polarization"

### Suppupporting Data.pdf

#### Supplementary Figure 1

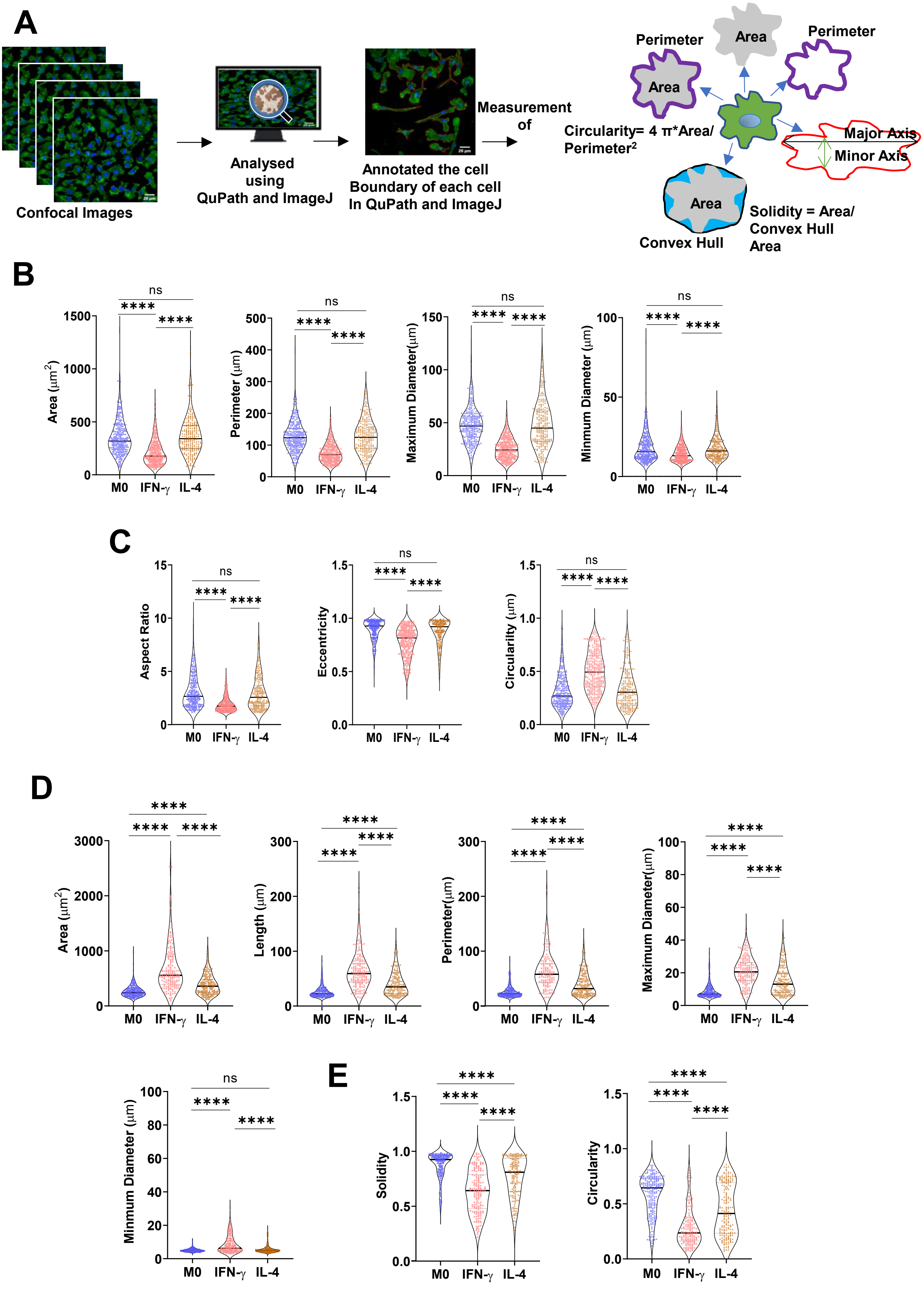

#### Supplementary Figure 2

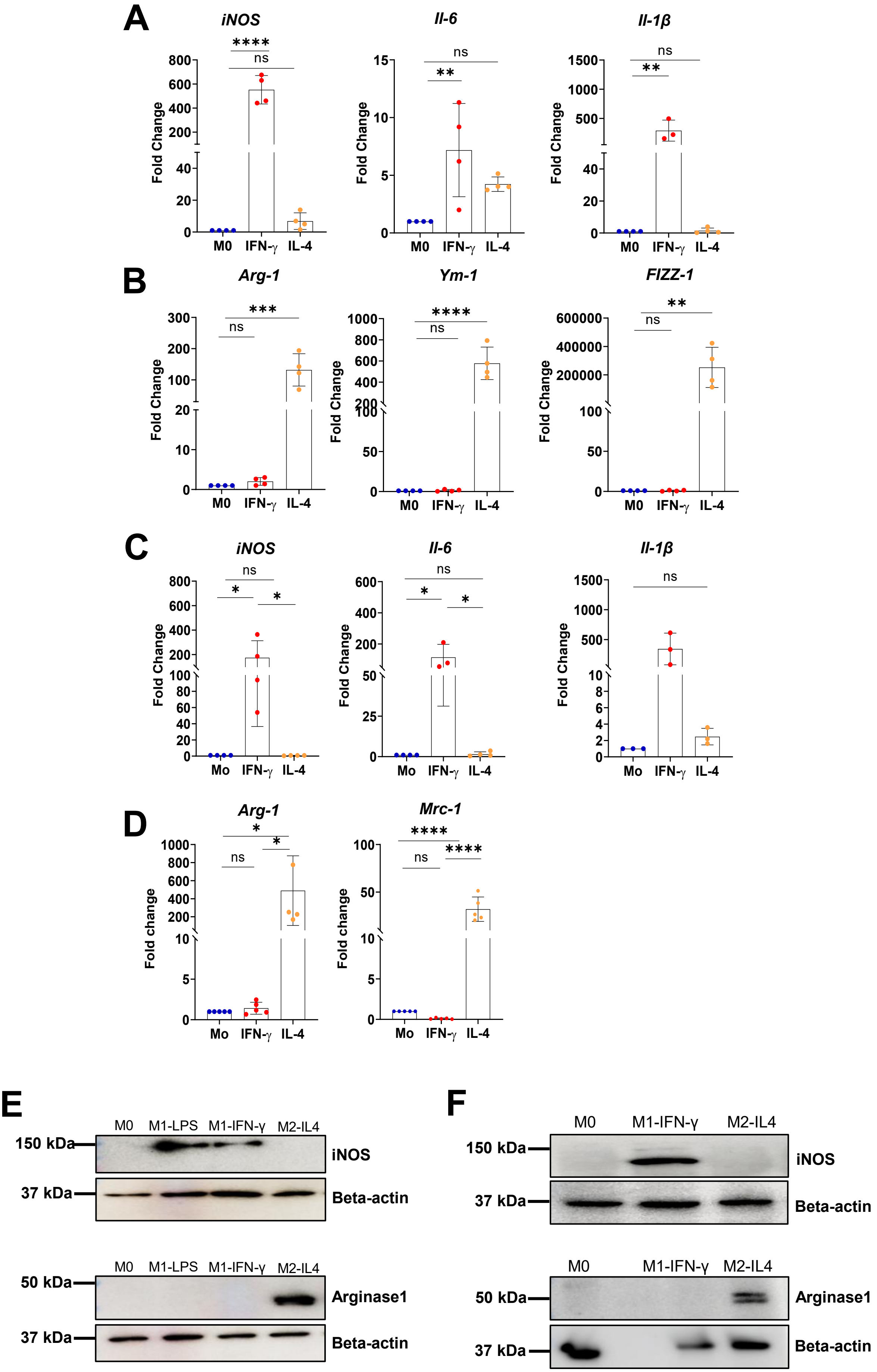

#### Supplementary Figure 3

## A

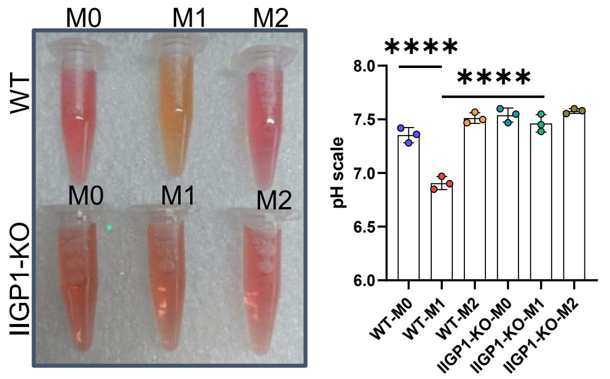
